# Cell type classification and discovery across diseases, technologies and tissues reveals conserved gene signatures and enables standardized single-cell readouts

**DOI:** 10.1101/2021.02.01.429207

**Authors:** Mathew Chamberlain, Richa Hanamsagar, Frank O. Nestle, Emanuele de Rinaldis, Virginia Savova

## Abstract

Autoimmune diseases are a major cause of mortality^1,2^. Current treatments often yield severe insult to host tissue. It is hypothesized that improved “precision medicine” therapies will target pathogenic cells selectively and thus reduce or eliminate severe side effects, and potentially induce robust immune tolerance^3^. However, it remains challenging to systematically identify which cellular phenotypes are present in cellular ensembles. Here, we present a novel machine learning approach, Signac, which uses neural networks trained with flow-sorted gene expression data to classify cellular phenotypes in single cell RNA-sequencing data. We demonstrate that Signac accurately classified single cell RNA-sequencing data across diseases, technologies, species and tissues. Then we applied Signac to identify known and novel immune-relevant precision medicine candidate drug targets (*n* = 12) in rheumatoid arthritis. A full release of this workflow can be found at our GitHub repository (https://github.com/mathewchamberlain/Signac).

## INTRODUCTION

The heterogeneity of autoimmune diseases complicates the discovery of new treatments. Similar symptoms can be associated with distinct immune phenotypes, and conversely, the same immune phenotypes can give rise to different symptoms. Rheumatoid arthritis (RA) is a prototypic example, where symptoms can vary and affect different organ systems not only across patients, but also longitudinally in the same patient^4,5^. Although knowledge of the immune phenotypes and their contributions to the disease profile would greatly enhance our ability to target pathogenic pathways in autoimmune diseases and in cancer, it is difficult to identify even a single cellular phenotype *in vivo*^6–8^, and therefore it remains unclear what is the immune composition of diseased tissues and how do cells contribute to the overall disease profile.

Due to their ability to identify cellular phenotypes in diseased tissue with single cell resolution, single-cell RNA sequencing (scRNA-seq) technologies are now mainstream in drug discovery and disease research^9^. However, cellular phenotypes are not identified from single cell data alone^10^. Instead, single-cell data analysis is typically a labor-intensive, project-specific and subjective process, such that two groups observing the same data will often arrive at a different set of conclusions^11^. The key analytical challenges of single-cell data analysis are how to map single-cell observations to known immune phenotypes in a consistent fashion independent of the scientist interpreting the experiment and how to efficiently identify disease-associated cellular phenotypes.

In this study, we filled this gap by developing a robust, efficient and scalable machine learning algorithm, Signac, which (a) accurately and consistently maps single-cell identities to a detailed hierarchy of known immune phenotypes; (b) identifies novel cell populations and (c) surfaces known and novel drug targets from single cell data. Overall, our approach converts scRNA-seq data into an objective readout that can be used for the study of immune cells across diseases, technologies, species and tissues^12^.

Our approach differs from all existing automatic annotation methods (**Table 1**) as it is the only method that; (a) reliably classifies single cell data from any technology or tissue without any tissue- or technology-specific training^11^; (b) was validated with CITE-seq and with flow cytometry data; (c) uses both single cell and bulk gene expression reference data, allowing it to “learn from experience”; (d) discovers potentially novel cellular populations in single cell data; (e) identifies rare cellular phenotypes (a known limitation of other methods)^13^; (f) differentiates cell types that are highly similar to each other, like T cell subsets (a known limitation of other methods)^11,13^; (g) classifies individual cells instead of clusters of cells^12,14^; (h) classifies non-human data to help study under-represented model organisms that lack sufficient reference data^11,13^; (i) is integrated with popular software packages SPRING and Seurat for ease of use^12,15^; and (j) outperformed all other pre-trained methods in benchmarking data from peripheral blood mononuclear cells (PBMCs) that were sequenced with seven different technologies (10X v2, 10X v3, CEL-Seq2, Drop-seq, InDrop, Smart-seq2 and Seq-well)^16^. We attribute these differences to the novelty of our method: a neural network-based hierarchical classification trained with bulk sorted reference data which does not require pre-defined marker genes and can learn from the single cell data that it classifies. To summarize, its detailed immunological classification and its extensive ability to learn and refine cell type representations make our algorithm unique among existing solutions.

**Table 1:** Comparison of cell type annotation methods.

| Name (version) | Method | Reference data source | Trained prior to inter-dataset classification<sup>11</sup> | Unknown cell types allowed | CITE-seq validated | FACs validated | Inter-dataset performance<sup>†</sup> |
| --- | --- | --- | --- | --- | --- | --- | --- |
| scVI<sup>17</sup> (0.3.0) | Neural network | Single cell | Yes | No | No | No | Poor |
| Cell-BLAST<sup>18</sup> (0.1.2) | Cell-to-cell similarity | Single cell | Yes | Yes | No | No | Poor |
| Garnett<sup>19</sup> (0.1.4) | Elastic net multinomial regression | Single cell | Yes | Yes | No | No | Poor |
| SCINA<sub>DE</sub><sup>20</sup> (1.1.0) | Bimodal distribution | Single cell | Yes | No | No | No | Good |
| SingleCellNet<sup>21</sup> (0.1.0) | Random forest | Single cell | Yes | No | No | No | Good |
| CHETAH<sup>22</sup> (0.99.5) | Correlation (hierarchy) | Single cell | Yes | Yes | No | No | Poor |
| scmapcell<sup>23</sup> (1.5.1) | KNN classification with cosine similarity | Single cell | Yes | Yes | No | No | Good |
| scID<sup>24</sup> (0.0.0.9000) | Fisher's linear discriminant | Single cell | Yes | Yes | No | No | Good |
| SCINA<sup>20</sup> pre-trained (1.1.0) | Bimodal distribution | Single cell | No | Yes | No | No | Poor |
| Moana | SVM with linear kernel | Bulk | No | No | No | No | Poor |
| Garnett pre-trained<sup>11,19</sup> (0.1.4) | Elastic net multinomial regression | Single cell | No | Yes | No | No | Poor |
| DigitalCellSorter<sup>25</sup> (e369a34) | Voting based on markers | Marker-based | No | No | No | No | Poor |
| Signac (2.0.7) | Neural network | Bulk and single cell | No | Yes | Yes | Yes | Good |

<sup>†</sup>Defined for all methods except Signac previously<sup>11</sup>; we used the same metric here (see Methods: Benchmarking Signac against other annotation methods; Supplemental dataset 1)

## RESULTS

### A novel approach for immune cell identification (Signac)

To annotate cellular phenotypes in single-cell transcriptomic data, we developed a novel approach, Signac, which used machine learning to classify each cell in unlabeled scRNA-seqdata according to a detailed hierarchy of immune phenotypes (**Fig. 1A-B**). Our approach is based on an ensemble of neural network classifiers that were trained on a reference dataset of gene expression profiles for purified, sorted cell types derived from flow-sorted cells generated by the Human Primary Cell Atlas (HPCA)^26^. First, we identified gene markers that distinguished each level of the cell type hierarchy (**Fig. 1A**) by performing differential gene expression analysis with the HPCA data and by performing meta-analysis of previously established gene markers (**Supplemental Figure 1**; see <u>Methods: Establishing the HPCA reference data gene markers fortraining Signac</u>; **Supplemental Dataset 2**)^26–28^. This established a set of gene markers, but it leftopen the question of how to use them to identify cells in scRNA-seq data. We reasoned that this task could be accomplished with machine learning^29^. However, the HPCA data contains as few as two samples for each sorted cell type population (**Supplemental Figure 1**), which is too few for machine learning methods that typically require hundreds or thousands of samples^26,30^. To address this problem, we bootstrapped the HPCA data and then used the bootstrapped data to train an ensemble of *n* = 100 neural network classifiers to make cell type classifications (**Fig. 1C**; see <u>Methods: Establishing a predictive model for cellular phenotypes using the HPCA reference data</u>; **Supplemental Figure 1**).

**Figure 1:**
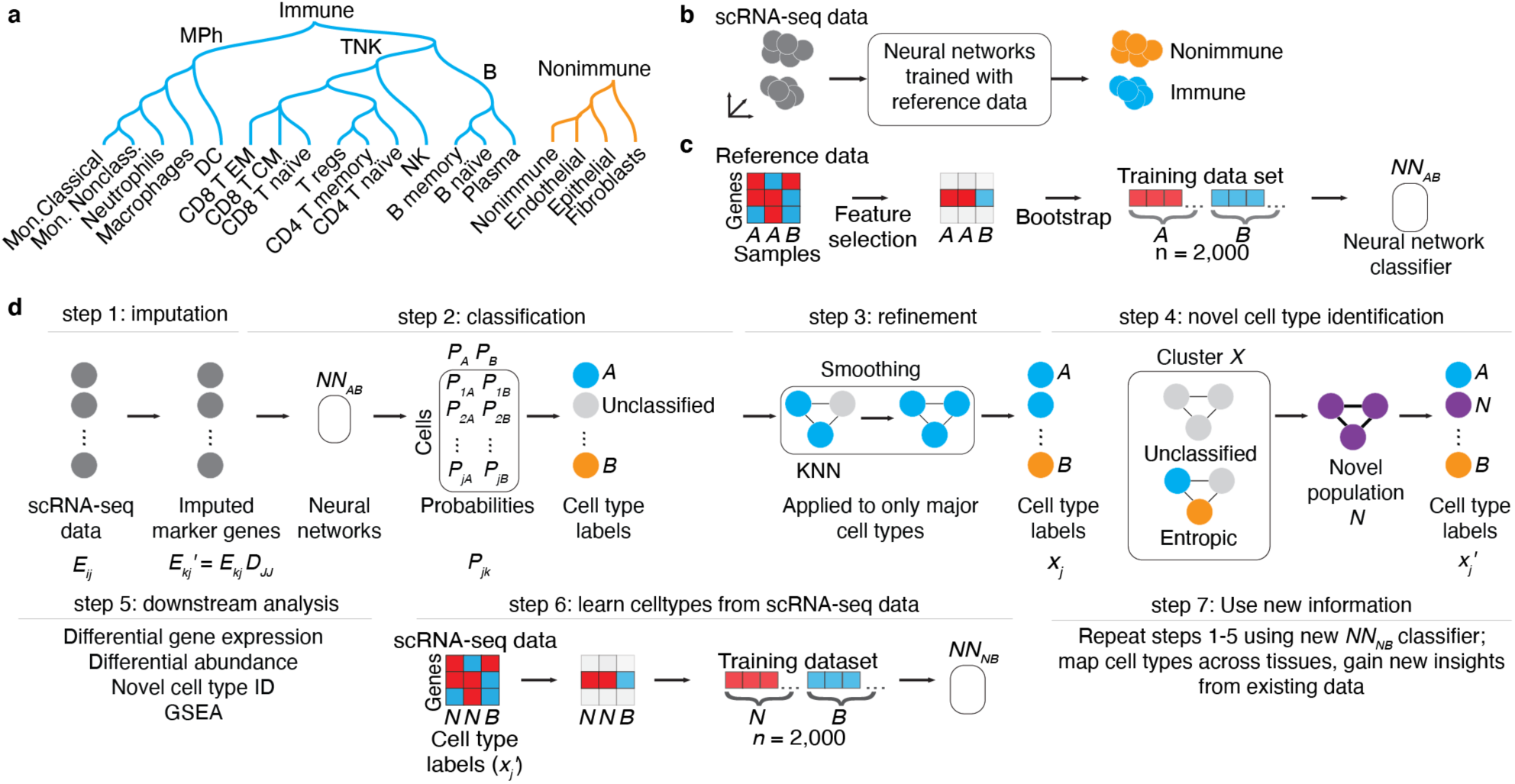
Conceptual overview of the Signac approach. **A, Classification hierarchy.** Dendrogram displays the hierarchy of cellular phenotypes that are classified by Signac: immune (teal) and nonimmune (carrot orange), major immune cell phenotypes (mononuclear phagocytes “MPh”; B cells and T/NK cells) and functional/terminally differentiated cellular phenotypes (rows). **B, Signac conceptual overview (theoretical data)**. Signac takes as input scRNA-seq data for which the cellular phenotype is unlabeled (left; *n* = 10 cell barcodes, grey circles). Next, Signac applies neural networks trained with pure, sorted reference data which results in labeled scRNA-seq data (teal; immune, carrot orange; nonimmune). The scRNA-seq data represented here are in a dimensionality-reduced plot (axes) where distances correspond to transcriptional similarities between cells (e.g., UMAP, t-SNE or PCA). **C, Concept for training neural network classifiers by bootstrapping a reference dataset (theoretical data)**. Heatmap (left) shows the expression (red indicates high gene expression, blue indicates low gene expression) of genes (*n* = 3, rows) across samples (*n* = 3, columns) in theoretical flow-sorted gene expression reference data of two pure cell type populations, *A* and *B*. Feature selection (black arrow) identifies a single gene that is correlated with *A* and *B*. This marker gene is bootstrapped (black arrow) by resampling from *A* and *B* separately, yielding a training data set for that gene, with a balanced number of bootstrapped samples from cell population A (*n* = 1,000 samples) and B (*n* = 1,000 samples). Next, neural network classifiers (*n* = 100) are trained (black arrow) on the training data set, yielding an ensemble of neural network classifiers (*NN*_*AB*_) that can be used to identify cell types *A* and *B*. **D, Example workflow (theoretical data)**. Signac takes as input scRNA-seq expression data (left; expression matrix *E*_*ij*_ with *i* = 1, …, *m* gene rows and *j* = 1, …, *n* cell columns) for which the cell type identity of each cell is unknown (gray circles). In step 1, a subset of the genes is imputed (arrow) using the imputation operator *D*_*jj*_ (see <u>Methods: KNN imputation</u>) yielding *E*_*kj*_′. Next, an ensemble (*n* = 100) of neural network classifiers (*NN*_*AB*_; black box) are applied to the imputed expression matrix *E*_*kj*_′, yielding for every cell a set of probabilities (one for each classifier) that the cellular phenotype is either phenotype *A* (*P*_*A*_) or *B* (*P*_*B*_). In step 2, these probabilities are then averaged and reported as a single probability, corresponding to the probability matrix *P*_*jk*_, and then each cell (circle) is amended a label (teal, carrot orange) corresponding to the maximal probability of *P*_*jk*_. Alternatively, a cell (circle) remains unclassified (light gray circle) if the maximal probability is below a user-set threshold. In step 3, after initial classification of cell types, KNN networks (black lines indicate network edges) are used to correct broad cell type assignments corresponding to immune and nonimmune cells and the first level of the cell type hierarchy (Fig 1A); each cell is assigned to the majority of itself and its first-degree neighbors in KNN networks. Classification continues until the deepest cell types in the hierarchy (Fig 1A), resulting in a vector of cell type labels *x*_*j*_. In step 4, novel cell types are identified using Louvain clustering to identify theoretical Cluster X (3 cells, black box); if cluster X is statistically enriched (two sample t-test, p-val < 0.01) for unclassified cells (top) or exhibits statistically significant normalized Shannon entropy in Signac labels (bottom), then Cluster X is flagged as a potential novel cellular population “N” (purple), yielding novel cell type labels (right). In step 5, the cell type labels are used for downstream analysis (listed). In step 6, scRNA-seq data is now used as a training data set to learn novel cell types (e.g., Novel population “N”) from scRNA-seq data, by developing neural network models that can distinguish novel cell populations (*N*) from other cell populations (*B*). Finally, this new model can be applied to other single cell data sets, yielding new classifications (step 7).

To validate our approach, we generated predictions for flow-sorted gene expression data that were not used in the analysis described above and instead originated from the Encode and Blueprint Epigenomics consortia, which used a different sequencing technology (RNA-seq) than the HPCA data (microarray)^26,31,32^. We observed 100% accuracy in the classification of B-cells, mononuclear phagocytes, neutrophils, CD8 T-cells, CD4 T-cells, NK-cells, plasma cells, T regulatory cells and nonimmune cells (**Supplemental Figure 2**; see <u>Methods: Signac classification</u>), supporting the idea that our approach can classify diverse cell types in different data sets^26^.

However, single cell data are distinct in many ways from the HPCA, Blueprint and Encode data described above^26,31,32^. For example, single cell data are sometimes composed of cell types for which flow-sorted data are unavailable, and may exhibit single cell technology-specific artifacts, like dropouts and doublets^10,27,33–35^. To address these concerns, we developed methods for learning gene expression-based representations of cell types from single cell data, for imputing missing gene expression values, and for leaving cells unclassified if they did not conform to a known cellular phenotype (**Fig. 1D**).

### Signac reliably distinguishes immune cells from non-immune cells in a variety of peripheral tissues

A fundamental requirement of automated immune cell type classification is the ability to distinguish immune cells from the cells of the host tissue at the infiltration site. Our approach succeeded in separating immune from nonimmune cells in data from three mixed tissue experiments deriving cells from human kidney, synovium, and lung, generated with either plate-based (**Fig. 2A-B**) or droplet-based technologies (**Fig. 2C**). These data were visualized with SPRING, a two-dimensional force-layout embedding that we used for interactive exploration of single-cell gene expression data (see <u>Methods: Single cell data pre-processing</u>)^15,36^. Signac also correctly rejected the non-immune label in data derived from human peripheral blood mononuclear cells (PBMCs; **Fig. 2D**)^37^, indicating accurate immune and nonimmune cell-classifications in peripheral tissues as well as in blood.

**Figure 2:**
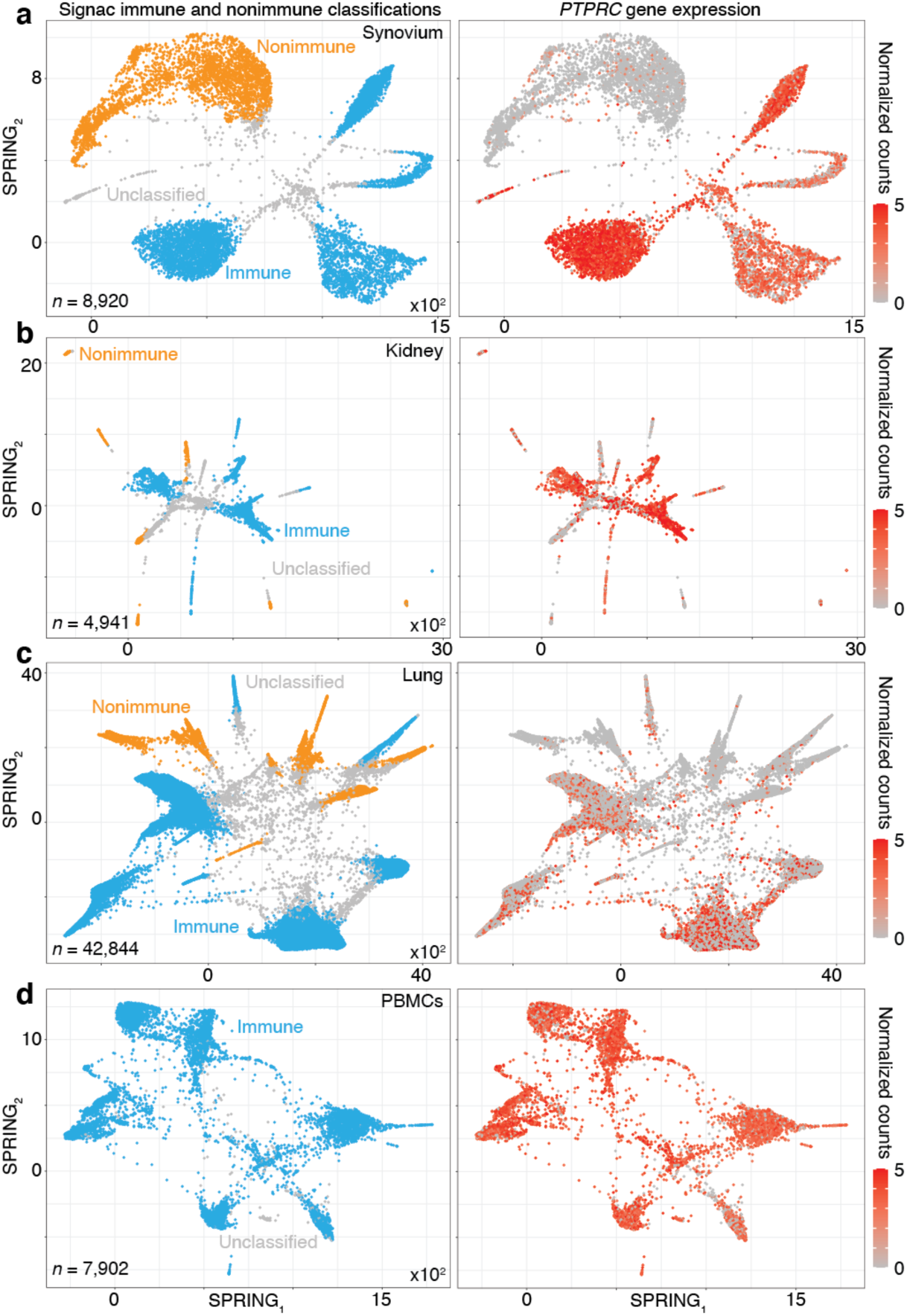
Signac reliably distinguishes immune and nonimmune cells in peripheral tissues. **A, Signac classifications were consistent with *PTPRC* expression in CEL-Seq2 data from synovium**. Two-dimensional visualization (left; SPRING plots) of single-cell transcriptomes (*n* = 8,920) in synovium biopsies (*n* = 26). Each cellular transcriptome (dot) was colored by Signac classifications; immune (teal), nonimmune (carrot orange) or unclassified (grey) cellular phenotypes. Single-cell gene expression plot (right) for a representative immune cell-type-enriched gene. **B, Signac classifications were consistent with *PTPRC* expression in CEL-Seq2 data from kidney**. See the caption for **Fig 2A**, except these visualizations correspond to single-cell transcriptomes (*n* = 4,941) from kidney biopsies (*n* = 36). **C, Signac classifications were consistent with *PTPRC* expression in 10X data from lung**. See the caption for **Fig 2A**, except these visualizations correspond to single-cell transcriptomes (*n* = 42,844) from lung biopsies (*n* = 18). **D, Signac correctly rejected the nonimmune labels in blood**. See the caption for **Fig 2A**, except these visualizations correspond to single-cell transcriptomes (*n* = 7,902) from PMBCs (*n* = 1).

### Signac accurately classified cell types in distinct tissues, technologies and diseases

Next, we applied Signac to annotate cellular phenotypes for the synovium and the PBMCs data introduced above. Unlike typical scRNA-seq data, these data contain simultaneous protein expression data for each individual cell measured with cellular indexing of transcriptomes and epitopes by sequencing (CITE-seq) for the PBMCs data and with flow cytometry for the synovium data^33,37^. To validate the cell type classifications generated by Signac, we determined to what extent Signac, which uses only transcriptional information, labeled cellular phenotypes that were consistent with the expected lineage-specific protein expression data.

Using only transcriptional data, Signac identified several distinct cellular phenotypes in PBMCs that were consistent with the expected protein expression data: CD19^+^ B-cells, CD19^+^CD25+ memory B-cells, CD19^+^CD25^-^*CCR7*^*+*^ naïve B-cells, CD14^++^CD16^-^ classical monocytes, CD14^+^CD16^++^ nonclassical monocytes, CD3^+^ T cells, CD45RA^+^CD4^+^ naïve T-cells, CD45RO^+^CD4^+^ T memory cells, CD4^+^TIGIT^+^*FOXP3*^*+*^ T regulatory cells, CD45RO^+^CD8^+^ T effector memory cells, CD56^+^CD3^-^ NK cells, *CLEC10A*^*+*^ dendritic cells (DCs), *MZB1*^*+*^ plasma cells and CD56^+^CD3^-^ NK cells (**Fig. 3A-C**; additional examples **Supplemental Figure 3**). Furthermore, well-known gene markers for these cell types were identified here with an unsupervised and unbiased analysis that identified immune <u>ma</u>rker <u>ge</u>ne<u>s</u> (IMAGES) from single cell data (see <u>Methods: Identifying IMAGES in scRNA-seq data</u>; **Fig. 3B**; **Supplemental Figure 4**).

**Figure 3:**
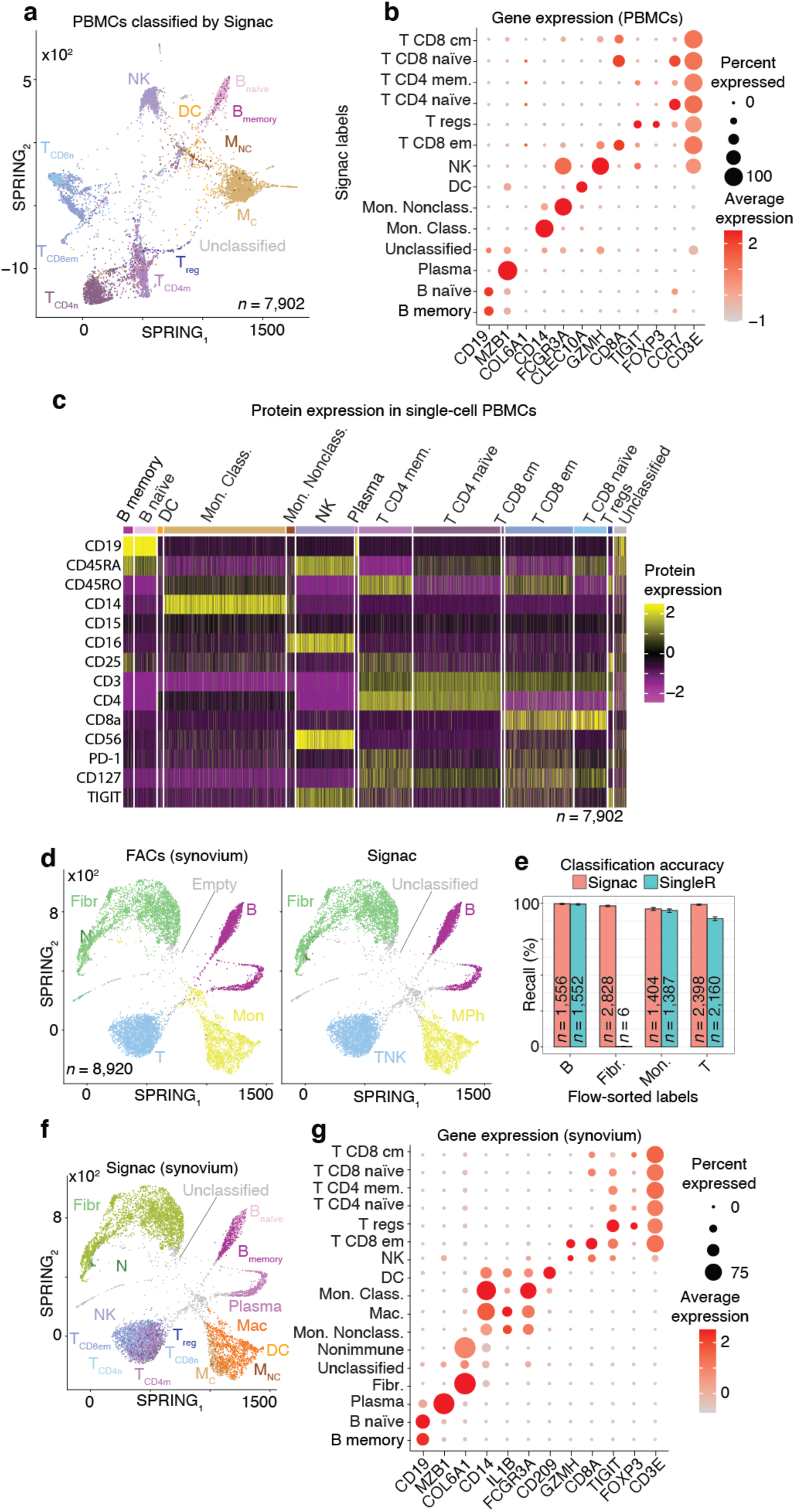
Validating Signac with single-cell protein expression data from PBMCs and synovium. **A, Two-dimensional visualization of Signac-classified CITE-seq PBMCs transcriptomes**. SPRING plot visualization as-depicted previously (**Fig 2D**) except with deeper Signac annotations for cell types. **B, Dot plot of top IMAGES expressed in CITE-seq PBMCs in cellular phenotypes labeled by Signac**. Dot plot shows the percentage (size) of single-cell transcriptomes within a cell type (y-axis) for which non-zero expression of marker genes was observed (x-axis). Color displays the average gene expression (red indicates more expression) in each cell type category. **C, Heatmap of protein expression in CITE-seq PBMCs in cellular phenotypes labeled by Signac**. Color shows the scaled protein expression data (rows’ yellow is higher expression; purple is lower expression) across single-cell transcriptomes (columns). Annotation bar indicates the cell type assigned by Signac (i.e., **Fig 3A-B**). **D, Two-dimensional visualization of synovium single-cell transcriptomes with cell types identified by FACs (left) and Signac (right)**. SPRING plot visualization as-depicted previously (**Fig 2A**) except with cell type labels determined by FACs (left), where each single-cell transcriptome is colored by the label assigned to it with flow cytometry (T cells, teal; fibroblasts, green; empty, grey; B cells, purple and monocytes, yellow). On the right, the same data are plotted the same way, except with labels generated with Signac. **E, Bar plot of Signac and SingleR performance in cell type classification with synovium**. Bar plot shows each flow-sorted cell type category (x-axis), and the performance of Signac (red) and SingleR (blue) in recalling the flow cytometry labels (error bars correspond to 95% confidence intervals, two-sided binomial test). **F, Two-dimensional visualization of synovium single-cell transcriptomes identified by Signac. G, Dot plot of top IMAGES expressed in single-cell transcriptomes from synovium in cellular phenotypes labeled by Signac**. See caption for **Fig. 3B**.

Although this demonstrated the accuracy of Signac in one tissue, it remained unclear to what extent Signac classified cells in other biological contexts. Since the immune composition of synovium is known to be distinct from that of blood, it was advantageous to next study data from the Accelerating Medicines Partnership (AMP), which isolated cells from human joint synovial tissues and performed flow cytometry in addition to scRNA-seq^7,33^. The proteins observed in this study were well-established lineage-specific markers for four distinct cell types: CD45^+^CD3^+^ T cells, CD45^+^CD3^-^CD19^+^ B cells, CD45^+^CD14^+^ monocytes and CD45^-^CD31^-^PDPN^+^ fibroblasts^33^, which allowed us to compare flow cytometry labels established previously to those generated by our approach (**Fig. 3D**)^33^. Signac, using only the transcriptional measurements for each cell, identified 98.2% of the flow cytometry labels (95% C.I. [98.0%; 98.5%], p-value < 0.001, two-sided binomial test, *n* = 8,334 cells). Encouraged by this result, we compared Signac to another cell type annotation tool, SingleR, which uses pairwise correlations between reference transcriptomes and single cell data to make cell type classifications^11,26,27^. Signac outperformed SingleR in every cell type category, but most notably fibroblasts, the only nonimmune cell type in the data (**Fig. 3E**; see <u>Methods: Comparing Signac to SingleR</u>)^26,27,38^. Furthermore, Signac outperformed SingleR at low sequencing depths in immune cell type classification, generating accurate classifications with as few as 200 unique genes detected per cell (95.2% average recall; 95% C.I. [76.2%; 99.9%], p-value < 0.001, two-sided binomial test; *n* = 21 cells; **Supplemental Figure 5**), demonstrating that Signac was robust and classified cell barcodes at low sequencing depths. Next, we turned our attention to the ability of Signac to classify cellular phenotypes that extended beyond the flow cytometry panel (**Fig. 3D**) to the deepest level of Signac annotations (**Fig. 1A**), resulting in new cell type annotations for the synovial cells (**Fig 3F**)^33^. To help validate these annotations, the IMAGES identified here were consistent with well-established gene markers for molecular phenotypes, like *FOXP3* in T regulatory cells (**Fig 3G**; **Supplemental Figure 6**)^28^, which suggested that Signac had made accurate cellular phenotype classifications. However, we note that *CD19* transcript was detected in only 46.9% (*n* = 734 / 1,564) of the flow-sorted CD45^+^CD3^-^CD19^+^ B cells, which demonstrates the importance of using more than one gene marker to identify cellular phenotypes in scRNA-seq data.

Altogether, this demonstrated that Signac accurately labeled cellular phenotypes in two distinct experiments deriving cells from either blood (**Fig. 3A**) or synovium (**Fig. 3E**), from either healthy (**Fig. 3A**) or diseased samples (**Fig. 3E**) and using either droplet-based (**Fig. 3A**) or well-based (**Fig. 3E**) technologies.

### Signac learned and reliably classified rare CD56^bright^ NK cells across tissues

Next, we challenged Signac to learn a gene expression-based representation of a rare cell type from single cell data. To explore this idea, we studied a cellular phenotype that is increasingly important in the study of autoimmune diseases and cancer, the CD56^bright^ NK cells^39–44^. To identify this population, we followed a strategy typically used in flow cytometry; we defined CD56^bright^ NK cells with CD16 and CD56 protein expression in the CITE-seq data from PBMCs described above (**Fig. 4A-B**)^39^. To help validate that these cells were CD56^bright^ NK cells, we noted that these cells (a) were identified using CD56 and CD16 protein expression data similar to flow cytometry^45^; (b) expressed known gene markers of CD56^bright^ NK cells, such as *CCL5*^-^, *GZMB*^-^*/H*^-^*/K*^+^, *KLRC1*^+^, *PRF1*^-^, *SELL*^+^ and *XCL1*^+^ that were identified here with an unbiased, unsupervised approach that compared NK cells to CD56^bright^ NK cells (**Fig. 4C;** *n* = 31 marker genes detected; see <u>Methods: Differential gene expression analysis</u>; **Supplemental Dataset 3**)^45–48^; (c) were a minority subset of the NK cells that were detected (*n* = 12 CD56^bright^ NK cells out of 939 NK cells; 1.3%; 95% C.I. [0.7%; 2.2%]; two-sided binomial test), consistent with the expected rarity of CD56^bright^ NK cells in human blood^45^; and (d) the marker genes (*n* = 31) identified here were statistically enriched for a chemokine receptor pathway that includes *CCL5, CCR7, XCL1* and *XCL2* (see <u>Methods: Gene set enrichment analysis</u>), which is a known functional molecular phenotype of CD56^bright^ NK cells in human blood^45^.

**Figure 4:**
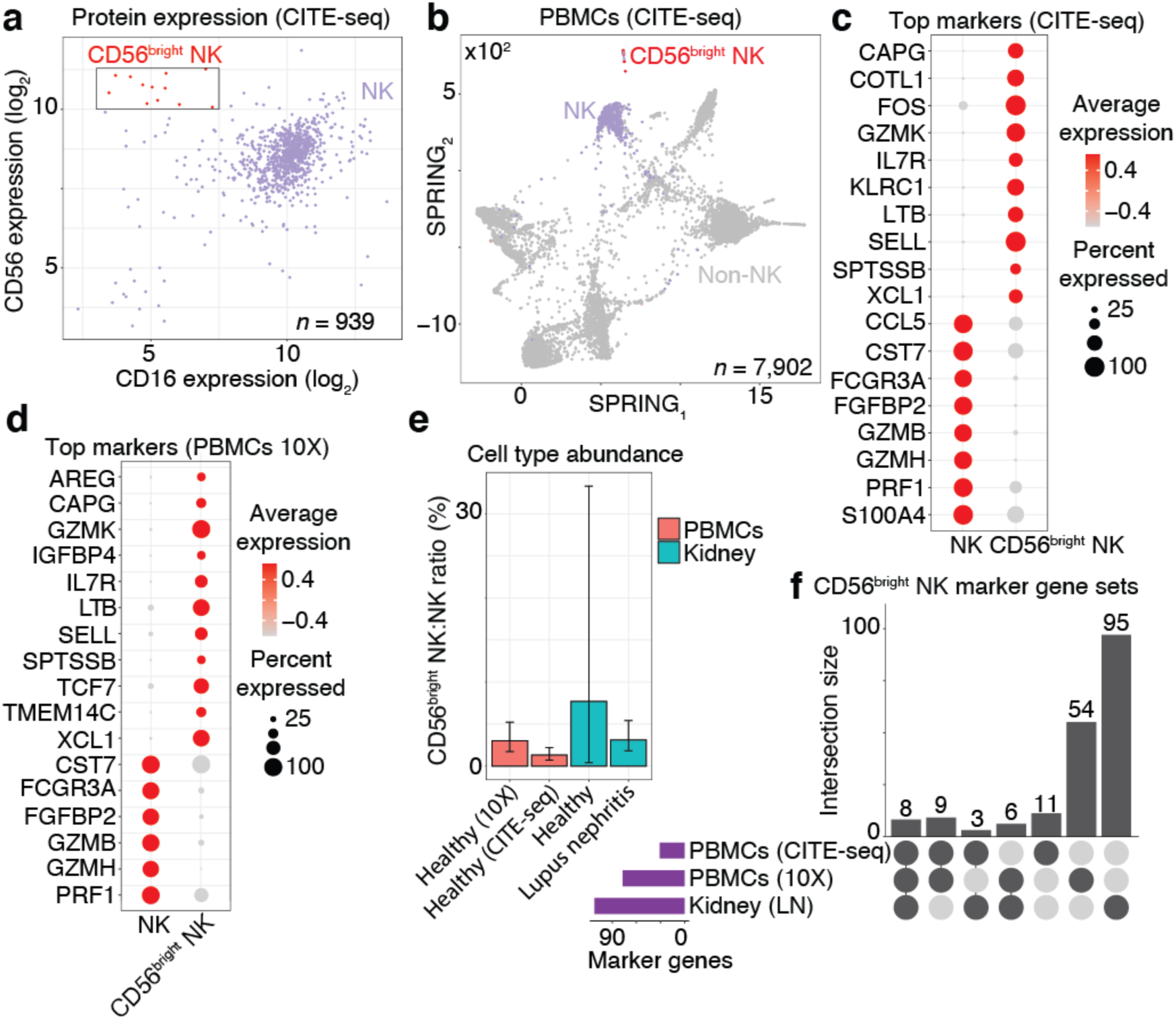
CD56^bright^ NK cells were learned from CITE-seq data and then classified in PBMCs and kidney data. **A, Scatter plot of protein expression in CITE-seq data revealed a population of CD56**^**bright**^ **NK cells (red; box)**. Scatter plot shows the CD56 and CD16 protein expression for NK cells (dots). **B, Two-dimensional visualization of the CITE-seq PBMCs single-cell data**. SPRING plot visualization as-depicted previously (**Fig. 3A**) except annotating just the NK cells (purple) and the sub-population of NK cells identified in **Fig. 4A** as CD56^bright^ NK cells (red). **C, Dot plot of top NK cell markers expressed in CITE-seq PBMCs**. Gene expression patterns across NK cell types; size of each dot indicates the percentage of single-cell transcriptomes within each cell populations (x-axis) for which non-zero gene expression (y-axis) was observed. Color displays the average gene expression (red indicates more expression) across single cell transcriptomes detected in each category. **D, Dot plot of top NK cell markers expressed in 10X PBMCs**. See the caption for **Fig 4C**, except these *n* = 17 marker genes were identified by classifying the CD56^bright^ NK in *n* = 4,784 single-cell transcriptomes from a different human sample and then performing differential expression analysis (resulting in *n* = 77 gene markers). The *n* = 17 plotted here were markers in both the CITE-seq and 10X data. **E, CD56**^**bright**^ **NK abundance bar plot in healthy blood, healthy kidney and lupus nephritis kidney**. Each bar is the ratio of single-cell transcriptomes classified as CD56^bright^ NK cells divided by all NK cells within each tissue (error bars are 95% C.I.; two-sided binomial test). These results were derived from *n* = 4,784 healthy PBMCs from one donor (10X), *n* = 7,902 healthy PBMCs from one donor (CITE-seq data described above), *n =* 501 healthy kidney cells from 8 biopsies, and *n* = 4,440 lupus nephritis kidney cells from 28 biopsies. **F, Upset plot reveals the number of NK cell markers that are shared across single cell data from blood and kidney**. Dark circles in the matrix (below) indicate sets that are part of the intersection. Bar plot (top) is ordered left-to-right by the largest intersecting set size; each number (top) indicates the number of marker genes belonging to that set. Bar plot (left) shows the number of marker genes identified in each data set (purple).

Next, we sought to identify these cells in tissues that lacked protein expression data by performing additional training with the CD56^bright^ NK cells serving as reference data for the Signac approach (**Fig 1C**; see Methods: Establishing a predictive model for CD56^bright^ NK cells <u>from CITE-seq PBMCs</u>). We used the “NK cell model” and classified additional single cell data from other tissues and technologies, which revealed that the CD56^bright^ NK cells were a conserved molecular phenotype that appeared with consistent abundance and with universal expression of eight marker genes across data derived from kidney and blood: *CAPG*^+^, *CST7*^-^, *FCGR3A*^-^ (CD16^-^), *FGFB2*^-^, *GZMB*^-^, *IL7R*^+^, *PRF1*^-^, *TCF7*^+^ (**Fig. 4D-F**). Altogether, this demonstrated that Signac learned a rare cell type from single cell data and then identified molecularly similar cells in other contexts^33,49^.

### Unclassified cells are mostly doublets; classified cells are mostly singlets

Next, we studied the behavior of Signac in the context of doublets, which are well-known artifacts in single cell data, by analyzing scRNA-seq data from human PBMCs that were analyzed previously in a study of *in silico* doublet detection using Scrublet (**Supplemental Figure 7**; see <u>Methods: Doublet detection in scRNA-seq PBMCs</u>)^50^. We found that every cellular barcode annotated as unclassified by Signac was classified as a doublet by Scrublet, whereas cells classified as either cell types or as novel cell type populations were mostly singlets, demonstrating that Signac accurately discerned known and novel cellular phenotypes from single cell artifacts.

### Signac identified conserved IMAGES across distinct tissues, technologies and diseases

Next, we turned our attention to characterizing the stunning diversity of cellular phenotypes in the human immune system. To help identify a universal molecular profile of immune cell phenotypes, we determined to what extent IMAGES were conserved in a trans-human study of single cell gene expression data across human donors (**Fig 5A**). To help validate the performance of Signac, we note that in contrast with data from kidney, lung and skin, Signac did not detect a single macrophage in PBMCs from any human sample, consistent with the idea that differentiation from monocytes occurs in tissue and not in blood (**Fig. 5B**; synovium data were excluded from this analysis because those cells were flow-sorted prior to sequencing)^33,51^. Next, we identified IMAGES for each human sample using the deepest Signac annotations (**Fig. 1A**), and then pooled them to identify IMAGES that were highly conserved (universal markers) across human samples and phenotypes, revealing known and novel gene markers (**Fig. 5C**; **Supplemental dataset 4**; see <u>Methods: Identifying IMAGES in scRNA-seq data</u>).

**Figure 5:**
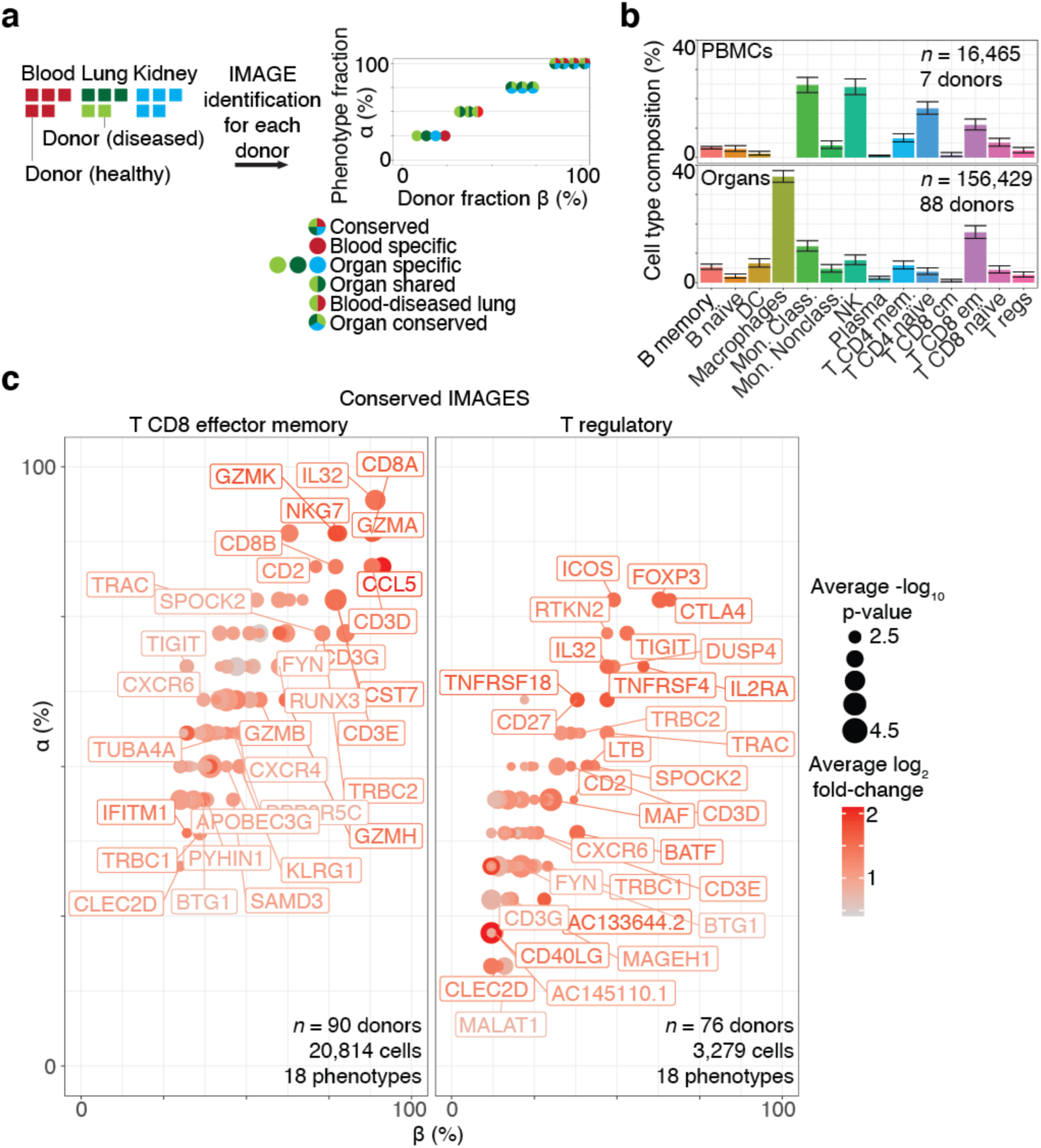
Systematic identification of conserved immune cell phenotypes with trans-human single cell gene expression study. **A, Overview of the approach (example workflow with theoretical data)**. Cells were extracted from humans (*n* = 15) representing four distinct biological phenotypes (colors: healthy blood, red; healthy lung, dark green; diseased lung, light green; kidney, teal), each from an individual human donor (*n* = 15). ScRNA-seq was performed for each donor individually followed by read mapping, normalization, filtering, immune cell classified by Signac, and then IMAGES were identified for each donor (arrow). Scatter plot displays the percentage of the four phenotypes (y-axis) and the percentage of the donors (x-axis) for which a gene (each dot is a unique gene) was identified as an IMAGE (colors indicate the phenotypes for which the gene was an IMAGE). Legend (right) shows the possible combinations. **B, Bar plot shows the average immune cell type composition of blood and organ samples classified by Signac**. The percentage of immune cells (y-axis) of each cellular phenotype (x-axis) classified by Signac. Results were average across donors; error bars were determined using the standard error of the mean. **C, Scatter plot revealed conserved IMAGES for T regulatory and T CD8 effector memory cells**. Scatter plot as depicted in **Fig 5A**. Each dot is a conserved IMAGE. Average log_2_ fold-change (colors) and p-values (size) were computed across human donors.

### Signac identified conserved and distinct gene expression patterns across species

Next, we challenged Signac to classify single cell data from model organisms for which flow-sorted datasets were generally lacking. We performed scRNA-seq of cynomolgus monkey PBMCs from three donors. Remarkably, Signac performed cell type classification without any species-specific training by mapping homologous gene symbols from monkey to human prior to classification (**Supplemental Figure 8**; see <u>Methods: Cross-species classification of single cell data from cynomolgus monkey PBMCs with human reference data</u>).

### Disease biology surfaced from single cell data with Signac

Next, we identified therapeutic opportunities for RA using single cell data. Together with clinicians, we hypothesized that the ideal treatment for RA would engage pathogenic immune cells precisely, and thereby prevent or reduce side effects and insult to host tissue, perhaps even eliminating the need for continuous treatment^3^. Although we have demonstrated that we can identify pathogenic immune phenotypes precisely with single cell data using Signac, finding a potential gene target was challenging because we lacked information about the expression of each gene in immune cells elsewhere in the body, risking the very off-target effects that we sought to avoid, and cross-tissue comparisons are notoriously difficult due to unintended technical artifacts (“batch” effects)^12,52^.

Here, we identified *n* = 24 genes as potential drug targets for RA on the basis that these genes were (a) in the initial pool of drug target candidates (see <u>Methods: Establishing an initial pool of drug target candidates</u>); (b) IMAGES for CD8^+^ effector memory T or naïve B cells in biopsies from RA synovium; and (c) not IMAGES for T regulatory cells in synovium (RA and OA), PBMCs (healthy and NSCLC), lung (NSCLC, sarcoidosis, ILD, IPF and healthy), kidney (lupus nephritis, renal carcinoma, healthy), or skin (healthy, atopic dermatitis lesions and non-lesions). Altogether, these genes were expressed specifically in pathogenic cellular phenotypes and not in T regulatory cells, and, compared to the initial pool of drug target candidates, the potential drug targets identified here were significantly enriched for therapeutic targets that were either in clinicals trials or FDA approved already for an immune condition, consistent with our expectations that robust immune cell phenotype classification surfaces immune-relevant therapeutic targets (**Figure 6A-B**)^3,33,36,53^.

**Figure 6:**
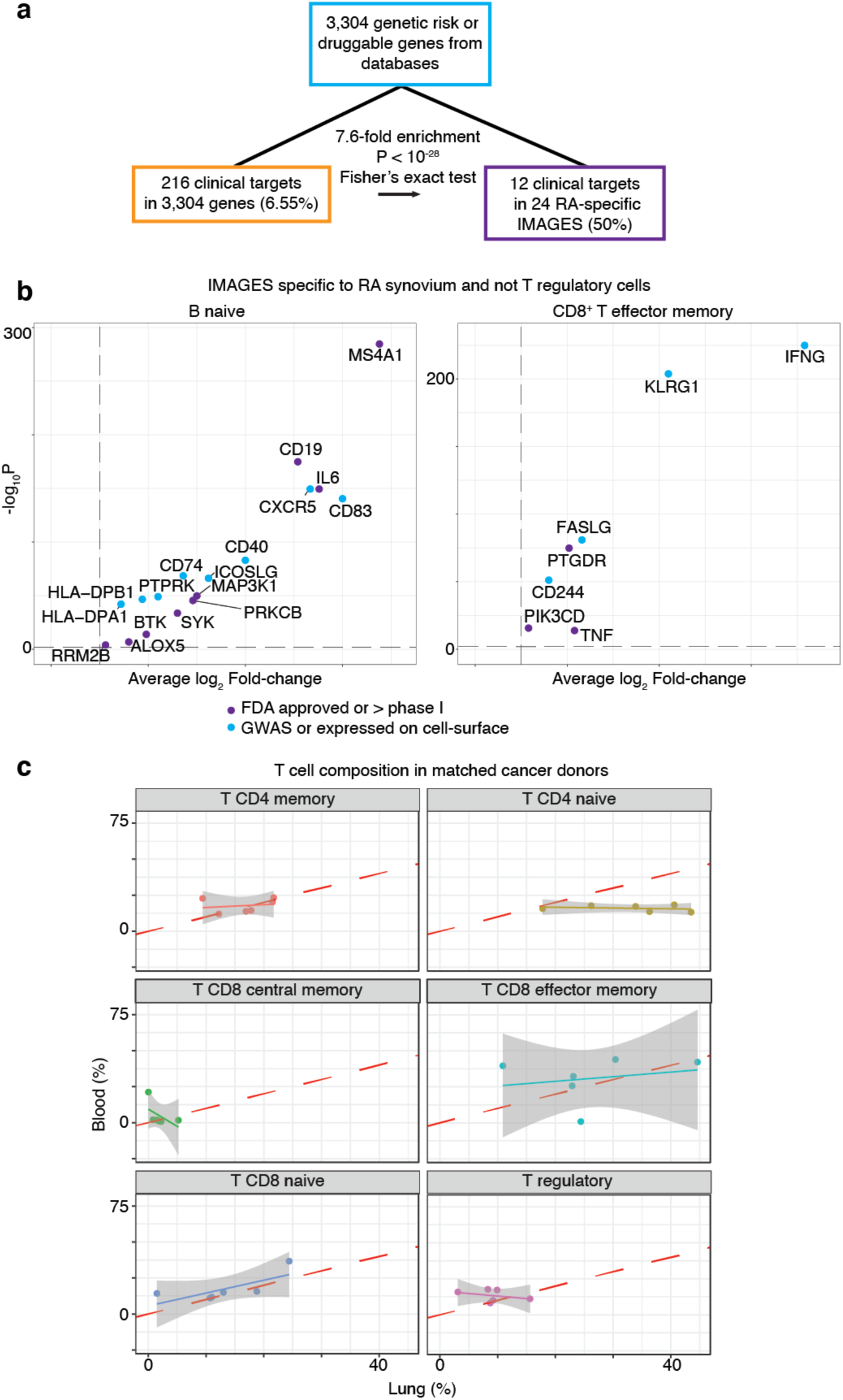
Disease biology surfaced from single cell data with Signac. **A, Overlap of drug targets and IMAGES identified here with enrichment score**. The initial set of genes (teal) contained *n* = 216 clinical targets in 3,304 genes (orange). We identified *n* = 24 RA-specific IMAGES (purple) and this set was enriched for clinical targets. **B, Volcano plot shows the IMAGES for the *n* = 24 potential target genes in RA**. Scatter plot shows the IMAGES identified here colored by clinical targets (purple) and genes that were in the initial set of genes, but not clinical targets (teal). **C, Scatter plot reveals the correlation between blood and lung composition in matched NSCLC samples**. Each dot is a donor for which a sample was taken from blood (y-axis) or lung (x-axis); axes correspond to the percentage of each phenotype among T cells classified by Signac.

Finally, we demonstrate that in matched samples taken from either blood and lung from humans diagnosed NSCLC, there was strong correlation in the composition of CD8^+^ T cells in blood and lung cancer, suggesting that blood-based biomarkers using these cellular phenotypes may someday supplant the need for lung cancer tissue biopsies (**Figure 6C**).

## DISCUSSION

The ability to uniformly and accurately identify known immune cell types in single cell data is a bottleneck for the application of single cell technology in pre-clinical and clinical studies of autoimmune disease and cancer. There are a number of technical challenges in the development of a cell type classification algorithm, stemming from the diversity of gene expression across tissues and diseases, the relative paucity of unique gene expression-based markers for each cell category and the high number of dropout measurements inherent to single cell transcriptomic data. Here, we demonstrated that our approach was robust to tissue, disease status, sequencing depth, sequencing technology and even performed well with closely related species for which training data was not readily available. Our approach was originally trained on transcriptional data from sorted bulk samples, but also used single cell data to refine representations of existing categories and learn new ones. Importantly, it also flagged potentially novel cell types for expert curation. However, the typical single cell experiment offers only a limited view of the human immune system as a typical experiment only analyzes the micro-environment of a single tissue, leaving unobserved the vast and rich complexity of immune cells in the body. To overcome this challenge, we analyzed immune cells in different biological contexts and in samples from different humans to reveal conserved and distinct IMAGES, and then we used this information to find potential immune-relevant precision drug targets from single cell data.

Current therapies for autoimmune diseases and cancer disrupt immune homeostasis, and sometimes give rise to new autoimmune syndromes or harmful immunosuppressive side effects^3^. Several new cell types have been identified recently with single cell technologies like CITE-seq, ATAC-seq and spatial transcriptomics. However, it remains unclear to what extent these cell types represent evidence of conserved cellular phenotypes that can be observed in other assays, or phenotypes that are unique to a given experimental protocol or observation method. As several high-profile projects strive to create an atlas of human cells, it is becoming increasingly important to learn gene expression-based cell type representations from these technologies to identify them when and where they appear in other assays.

In a typical workflow that studies the molecular identity of cells, one typically compares populations or clusters of cells, in which cells derived from distinct human samples are pooled together to form a group. Although this is useful for identifying broad patterns of similar cells, this also diminishes or eliminates the detection of sample heterogeneity (i.e., patient stratification), in which subsets of patient cells are enriched for a particular molecular identity. Several technologies, like gene expression-based biomarkers and cell type deconvolution algorithms like CIBERSORT, require well-established gene expression-based signatures for cell types, and thus it behooved us to identify gene signatures by studying individual samples first, and then pooling the results to identify conserved and distinct gene signatures, which we performed in **Figure 5**. Notable, all cell types were identified with the same unsupervised approach described above (Signac) without any changes to parameters or special considerations for any individual tissue or sample.

It is conventionally thought that machine learning methods require similar data types to train and to classify data. Here, we trained our models with data from microarray experiments with cellular ensembles and then used these models to classify single cell data from diverse tissues, accurately classifying synovial fibroblasts in single cell data despite using human foreskin fibroblasts in the HPCA dataset^26^. We believe that this work opens the door to a new wave of machine learning approaches that will integrate disparate data and create more uniform and complete pictures of cell biology.

## METHODS

### Benchmarking Signac against other annotation methods

To benchmark Signac against other annotation methods, we accessed the “PbmcBench” data – a resource of 19,792 human PBMCs sequenced across seven different technologies with cell type labels generated for each cell previously – from https://doi.org/10.5281/zenodo.3357167 on February 5, 2021^11^. Next, we classified scRNA-seq data from each of the seven technologies with Signac (v2.0.7) in R with the default parameters. Median F1 scores were computed as described previously^11^. Good inter-dataset classification performance (**Table 1**) was defined as having an average median F1-score > 0.75, as described previously^11^.

### Establishing the HPCA reference data gene markers for training Signac

To establish a reference dataset, we used the HPCA consortium data^26^, which comprised of 713 microarray samples annotated to 157 cell types, processed as described previously^27^ except that all genes that encoded for ribosomal proteins and mitochondrial transcripts were removed, all samples derived from bone marrow biopsies were removed, and we used a subset of genes that were identified as exhibiting cell type-specific gene expression previously (what remained was *n* = 10,808 genes and *n* = 544 samples corresponding to 113 annotated cellular phenotypes)^28^. The data as well as the cell type annotations for the HPCA reference data were accessed in R from the SingleR R package (v0.2.0)^27^.

To establish a set of gene markers for cell types using these data, we performed differential gene expression analysis comparing samples annotated as different cell types according to the cell type hierarchy (**Fig 1A**) and identified *n* = 5,620 genes that were significantly (p-value < 0.05, Wilcoxon-rank sum test; log-fold change > 0.25) differentially expressed using the Seurat package (v3.2.0) in R with the default settings, except that we used relative-count normalization instead of log normalization. This approach yielded no significantly differentially expressed genes for comparisons between memory and naïve B cells, plasma cells and B cells, memory and naïve CD4 T cells, T regulatory cells and CD4 memory T cells, memory and naïve CD8 T cells, and effector memory CD8 T cells and central memory CD8 T cells; in these cases we used *n* = 1,171 genes identified previously, which we accessed with the xCell package (v1.1.0) in R^27^. Altogether, the marker genes used here are available in **Supplemental Dataset 2**.

### Establishing a predictive model for cellular phenotypes using the HPCA reference data

To establish a predictive model for cellular phenotypes, the HPCA reference data (processed as described above) were split into disjoint subsets according to the cell type hierarchy (**Fig. 1A**; **Supplemental Figure 1**). At each level of the hierarchy, the marker genes were bootstrapped by random resampling with replacement across samples annotated as belonging to the same cell type, resulting in *n* = 1,000 bootstrapped samples of each marker gene in each cell type population. We introduced noise to each bootstrapped marker gene by sampling from a random normal distribution with mean and standard deviation set by the mean and standard deviation of the bootstrapped genes, and then we performed max-min normalization across genes.

Next, we constructed *n* = 100 neural networks in R using the neuralnet package (v1.44.2) with the neuralnet function with the default settings except that the linear.output parameter which was set to false. Neural networks were trained with the bootstrapped reference training data after taking the intersection of the genes in the training data and the target data^54^. Neural network hyperparameters were validated using the caret package (v6.0-86) in R for immune and nonimmune cell type classification, which yielded the default neuralnet settings, which were subsequently used for all neural network models.

### Signac classification

Cell type labels were generated according to the maximal probability derived from the average of an ensemble of neural networks (*n* = 100) trained with the HPCA reference data as described above. For single cell data, any individual cell barcode was labeled “Unclassified” when it exhibited large (2 standard deviations greater than the mean) normalized Shannon entropy within four nearest neighbors of the KNN network computed with immune and main cell type labels (**Fig 1A**). A user-set threshold was introduced such that any cell barcode with maximal average probability less than the threshold were labeled “Unclassified” – herein this threshold was not used (set to zero). In single cell data, any cell barcodes labeled “Unclassified” that significantly (*p* < 0.01, hypergeometric test) populated a Louvain cluster were amended a “potential novel cell type” label (**Supplemental Figure 7**).

### K-nearest neighbor smoothing

To reduce classification errors of cell barcodes labeled by Signac, we constructed k-nearest neighbor (KNN) graphs as described previously^15^, and after classification of immune, nonimmune and major cell types as described above (See <u>Methods: Signac classification</u>), the broad label for each cell barcode was assigned to the most frequent label of itself and of the nearest neighbors for immune cell type and major cell types (**Fig 1A**).

### Single cell data pre-processing

Unless stated otherwise, all scRNA-seq data analyzed here started from unfiltered count data. First, we removed all cell barcodes that expressed fewer than 200 unique genes and fewer than 500 counts. Next, we removed all cell barcodes with abundant (greater than 20% of the single cell library size) mitochondrial gene expression. Within this subset of cell barcodes, we removed all genes with zero detected counts, we removed all genes that were encoded for mitochondrial and ribosomal transcripts, and then library sizes were normalized to the mean library size of all cell barcodes. This procedure resulted in *n* = 8,920 cells in the synovium (**Fig 2A**), *n* = 4,941 cells in the kidney (**Fig 2B**), *n* = 42,844 cells in the lung (**Fig 2C**) and *n* = 7,902 cells in the CITE-seq PBMCs (**Fig 2D**). In the case of CITE-seq data (**Fig 2D**; **Fig 3A**), these same steps were performed only after setting aside the protein expression data. Protein expression data from CITE-seq were normalized with CLR normalization in R using the Seurat package (v3.2.0)^12^. Generation of a two-dimensional force-layout embedding was performed as described previously in Python with Jupyter notebooks that are available on our web-server^15^.

### Establishing an initial pool of drug target candidates

To establish an initial set of genes, we limited our analysis to genes that were druggable, associated with genetic evidence, or already approved by the FDA for an immunological condition as an established immune-relevant gene target. We accessed genes associated with genetic evidence from the GWAS catalog (version 1.0_e98_r2020-03-08) for any of the following immune conditions: rheumatoid arthritis, psoriatic arthritis, ankylosing spondylitis, giant cell arteritis, sarcoidosis, psoriasis, vitiligo, Crohn’s disease, ulcerative colitis, systemic lupus erythematosus, cutaneous lupus erythematosus, lupus nephritis in systemic lupus erythematosus", Sjögren’s syndrome, idiopathic pulmonary fibrosis, limited cutaneous systemic scleroderma, type 1 diabetes, celiac disease, asthma, chronic obstructive pulmonary disease, chronic rhinosinusitis with nasal polyps, atopic dermatitis, eosinophilic esophagitis, and peanut allergy. This yielded *n* = 2,326 unique genes. Next, we accessed all genes associated with genetic evidence and immune-relevant genes in clinical trials or approved by the FDA as those identified previously H. Fang et al., yielding *n* = 720 and *n* = 216 genes, respectively^55^. We identified genes expressed on the cell surface as those annotated as being a receptor, transmembrane protein, exhibiting peripheral expression, secreted or integrin with CellPhoneDB accessing the “protein_cureated.csv,” yielding *n* = 971 genes^56^. All total, this yielded *n* = 3,304 genes, which are provided with annotations in the Signac R package (v2.0.7).

### Establishing a predictive model for CD56^bright^ NK cells from CITE-seq PBMCs

To learn a gene-expression based representation of CD56^bright^ NK cells from single cell data, we first identified used CITE-seq to identify these cells with protein expression data (**Fig 4A-B**). Second, we defined a set of gene markers for the CD56^bright^ NK cells (*p*-value < 0.05, Wilcoxon-rank sum test; log-fold change > 1) with differential gene expression analysis that compared the CD56^bright^ NK cells to the non-CD56^bright^ NK cells in the CITE-seq data. Differential gene expression analysis was performed with the Seurat package (v3.2.0) in R using the FindMarkers function with the default settings which resulted in *n* = 31 marker genes. Third, we took a subset of the CITE-seq expression matrix corresponding to the *n* = 31 marker genes, performed KNN imputation (**Fig. 1D**), and then bootstrapped the single cell data as described above (**Fig 1C**). Lastly, neural network model training and subsequent classification was performed as described above (**Fig. 1**), except now each cell classified as “NK” using the HPCA reference data were further classified as CD56^bright^ NK cells and non-CD56^bright^ NK cells with the new neural network models. This workflow was executed by the SignacLearn function in R with the Signac (v2.0.7) package.

### Cross-species classification of single cell data from cynomolgus monkey PBMCs with human reference data

As described previously in our single-cell optimization study^9^, cryopreserved monkey PBMCs were thawed (2 vials at a time) in a 37°C water bath for 1-2 minutes until a small crystal remained. Cryovial was removed from the water bath and cell solution was transferred to a fresh, sterile 2 ml Eppendorf tube using a wide bore pipet tip. The cryovial was washed with 0.04% BSA/PBS and the solution was transferred to the Eppendorf tube. Sample was centrifuged at 150 rcf, 5 min, at room temperature (RT). Supernatant was carefully removed, and sample was washed with 1 ml of 0.04% BSA/PBS using wide bore pipet tip. Sample was re-centrifuged using the same conditions mentioned above. The cells were washed one more time for a total of 3 washes. After the final wash, cells were resuspended in 1 ml of 0.04% BSA/PBS and counted using manual hemacytometer and trypan blue. If the viability was found to be lower than 75%, the sample was subjected to a “clean-up” step using Dead Cell Removal kit (Miltenyi Biotec, Catalog #130-090-101). Cells were washed again and resuspended in 500 ul of 0.04% BSA/PBS and counted. Volume was adjusted to 1 million cells per ml of 0.04% BSA/PBS solution. After the cell volume was adjusted to 1 million per ml (or 1000 cells per ul), protocol for 10X Genomics 5’ v1 gene expression library preparation was used. 10,000 cells were targeted per sample. Quality of uniquely-indexed libraries was determined on the 2100 Bioanalyzer instrument (Agilent) using High Sensitivity DNA kit (Agilent, Catalog # 5067–4626) and quantified using Kapa library quantification kit (Kapa Biosystems, Catalog # KK4824 – 07960140001) on the QuantStudio 7 Flex Real-Time PCR system. The libraries were diluted in 10 mM Tris-HCl buffer and pooled in equimolar concentration (2 nM) for sequencing. Sequencing was performed on Nextseq2000. Sequencing depth and cycle number was as per 10X Genomics recommendations: Read 1 = 26 cycles, i7 index = 8 cycles, Read 2 = 98 cycles, and we aimed for a sequencing depth of 35,000 reads per cell.

Reads were aligned to the cynomolgus monkey genome which was built from the fasta file for M. *fascicularis* (v5.0) with the CellRanger (v3.1.0) mkref command with the default settings. After mapping, the “raw_feature_bc_matrix.h5” files generated by CellRanger were used for subsequent analysis in R. To map gene annotations, the M. *fascicularis* gene symbols were mapped to human gene homologs using annotations from the 2019 ensemble archive of the M. *fascicularis* genome and the 2019 ensemble archive version of the homo sapiens genome with the getLDS function in biomaRt (v2.38.0) in R. Next, we used a subset of the data corresponding to only counts mapped to genes that had a homologous human gene pair (*n* = 17,365 genes remained). When multiple M. *fascicularis* genes were homologous to a single human gene, any counts mapped to those genes were summed and reported as a single mapped gene with the homologous human gene symbol, resulting in *n* = 16,854 unique genes. Each cell barcode was then filtered as described previously (See Methods: <u>Single cell data pre-processing</u>), and then classified with Signac (v2.0.7) in R using the Signac function with the default settings.

### Comparing Signac to SingleR

To compare Signac to SingleR, we used the SingleR package (v0.2.0) in R and classified the synovium data with the SingleR function with the default settings, with the “ref_data” parameter set to the HPCA reference data attached to the SingleR package (**Fig 2D-E**; **Supplemental Figure 5**)^27^. We compared these results to Signac; we used the Signac package (v2.0.7) in R with the Signac function with the default settings.

### Differential gene expression analysis

Unless stated otherwise, differential expression analysis was performed with the Wilcoxon rank-sum test with an adjusted *p*-value cutoff (< 0.05), and a log-fold change cutoff (> 1) in R with the Seurat package (v3.0.2) in R using the FindMarkers function.

### Gene set enrichment analysis

Gene set enrichment analysis was performed in R with the ReactomePA package (v1.26.0) in R using the enrichPathway function with the default settings with thresholds of 0.05 to the adjusted p-value and 0.05 to the FDR^57^.

### Identifying IMAGES in scRNA-seq data

To identify immune marker genes (IMAGES) in single cell data, we identified genes that were significantly differentially expressed (p-value < 0.05, Wilcoxon-rank sum test; log-fold change > 0.25) in a “one verse all” comparison only among single cell transcriptomes that were annotated as immune cell phenotypes by Signac (v2.0.7) in R. Differential expression testing was performed with the Seurat package (v3.2.0) in R using the default settings.

To identify conserved IMAGES (**Fig. 5**), we performed the IMAGE analysis described above, except that it was performed within each human sample that had at least *n* = 200 detected immune cells, resulting in *n* = 178,929 immune cell barcodes across *n* = 114 human samples deriving cells from *n* = 18 distinct disease-tissue phenotypes corresponding to PBMCs (healthy, four stages of NSCLC), kidney (healthy, lupus nephritis and renal carcinoma), lung (healthy, IPF, four stages of NSCLC), and skin (heathy, atopic dermatitis from lesions and non-lesions) biopsies; all from previously published single cell studies^33,36,58–61^. The IMAGES for each human sample were pooled together, and then we reported the top genes (*n* = 100; **Fig. 5C**) for each cellular phenotype corresponding to the most fractional appearance of each gene as an IMAGE across distinct human donors.

### Doublet detection in scRNA-seq PBMCs

To classify doublets in PBMCs, we used Scrublet (v0.2.1) in python with the Scrublet and scrub_doublets functions with parameters set previously in the orginal Scrublet study^50^. The scRNA-seq data from PBMCS were accessed from 10X genomics (https://support.10xgenomics.com/single-cell-gene-expression/datasets/2.1.0/pbmc8k). To generate visualizations (**Supplemental Figure 7**), we used Seurat v3.2.0 in R with the default settings for each function: CreateSeuratObject, NormalizeData, FindVariableFeatures, ScaleData, RunPCA, FindNeighbors, FindClusters, RunUMAP and FindMarkers. To annotate cellular phenotypes (**Supplemental Figure 7**), we used Signac (v2.0.7) in R with the Signac function with the default settings, which was applied directly to the Seurat object using KNN edges identified with the FindNeighbors function.

### KNN Imputation

To impute gene expression values, the total number of genes detected in each cell was set to the diagonal of a cell-by-cell matrix *W*_*jj*_. Next, we established cells with direct and higher *k*-degree connections in the KNN network from the adjacency matrix *A*_*jj*_ and from K^th^ powers of *A*_*jj*_, forming a KNN network-based imputation operator *D*_*jj*_ which was weighted by the total number of genes detected in each cell, and normalized such that each row sums to two:

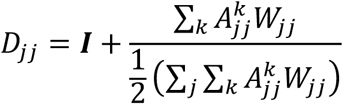

The imputed expression matrix 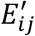 is then computed directly by operating on the observed expression matrix *E*_*ij*_. Here, we set *k* = 1 to use gene expression values within first nearest neighbors in the KNN network, resulting in the imputed gene expression matrix:

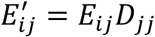

### Data and software availability

All data reported here are publicly available. The kidney and the synovium (**Fig. 2**) datasets were downloaded via ImmPort (accession codes SDY997 and SDY998, April 2019 release) from the AMP consortium^33^. The PBMCs CITE-seq data (**Fig. 3**) and healthy control data (**Fig. 5**) were downloaded from the 10X website (https://support.10xgenomics.com/single-cell-gene-expression/datasets/3.0.2/5k_pbmc_protein_v3 and https://support.10xgenomics.com/single-cell-gene-expression/datasets/3.0.2/5k_pbmc_v3). The blood (**Fig. 6**) and lung (**Fig. 6**) NSCLC data sets were downloaded from the NCBI GEO depository (accession number GSE127465)^36^. All software used in this study is available on the GitHub page for Signac (https://github.com/mathewchamberlain/Signac).

## Acknowledgments

We would like to thank members of the de Rinaldis lab for insightful discussion: H.X., R.M., Z.D., B.S., L.Z., C.R., D.M., P.T., M.S., P.C., V.A., and A.B. We would like to thank Reza Olfati-Saber for revisions to the manuscript. This work was supported by Sanofi US.

## Author contributions

M.C., E.D.R., V.S. and F.N. designed the research; M.C. performed the research, developed Signac and performed computational analysis of data with the guidance of V.S.; R.H. performed single-cell experiments with the guidance of V.S.; M.C., R.H. and V.S. drafted the manuscript and all authors contributed to writing the final version.

---

**Supplemental Figure 1:**
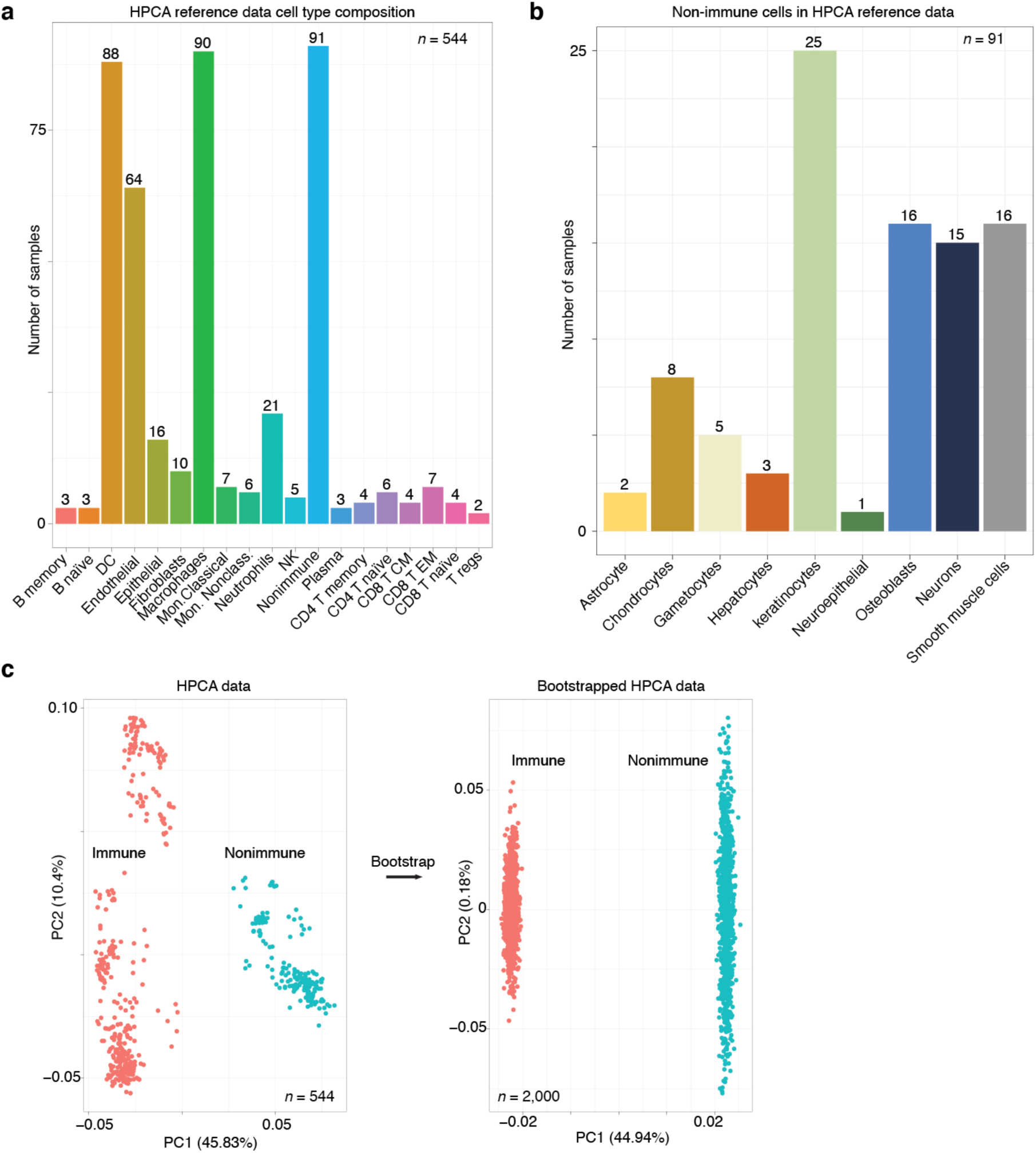
Overview of the HPCA reference data. **A, Number of samples bar plot for each annotated cell type**. Bar plot indicates the number of samples (y-axis; numbers) for each cellular phenotype (x-axis) that was in the HPCA reference data set. **B, Number of samples bar plot for the “nonimmune” cellular phenotypes**. Bar plot as depicted in **Supplemental Figure 1A** for the *n* = 91 samples in the “nonimmune” cell type category. **C, Principal component analysis (PCA) plots revealed that the HPCA reference data and the bootstrapped training data were separable by immune and nonimmune gene markers**. PCA plot (left) shows a gene expression sample (each dot) from the HPCA reference data that is either labeled as immune (red) or nonimmune (teal) based on the classification hierarchy established here and the annotated cell type established experimentally (**Fig. 1A**). PCA was performed on the marker gene that were determined by differential gene expression analysis comparing the samples annotated as immune and nonimmune. After bootstrapping (arrow); PCA was performed with the same marker genes, except with data that was bootstrapped from the immune and nonimmune samples. We note that the structure in the PCA plot prior to bootstrapping was largely removed, consistent with the view that bootstrapping generated what can be thought of as composite or as average immune and nonimmune gene expression samples.

**Supplemental Figure 2:**
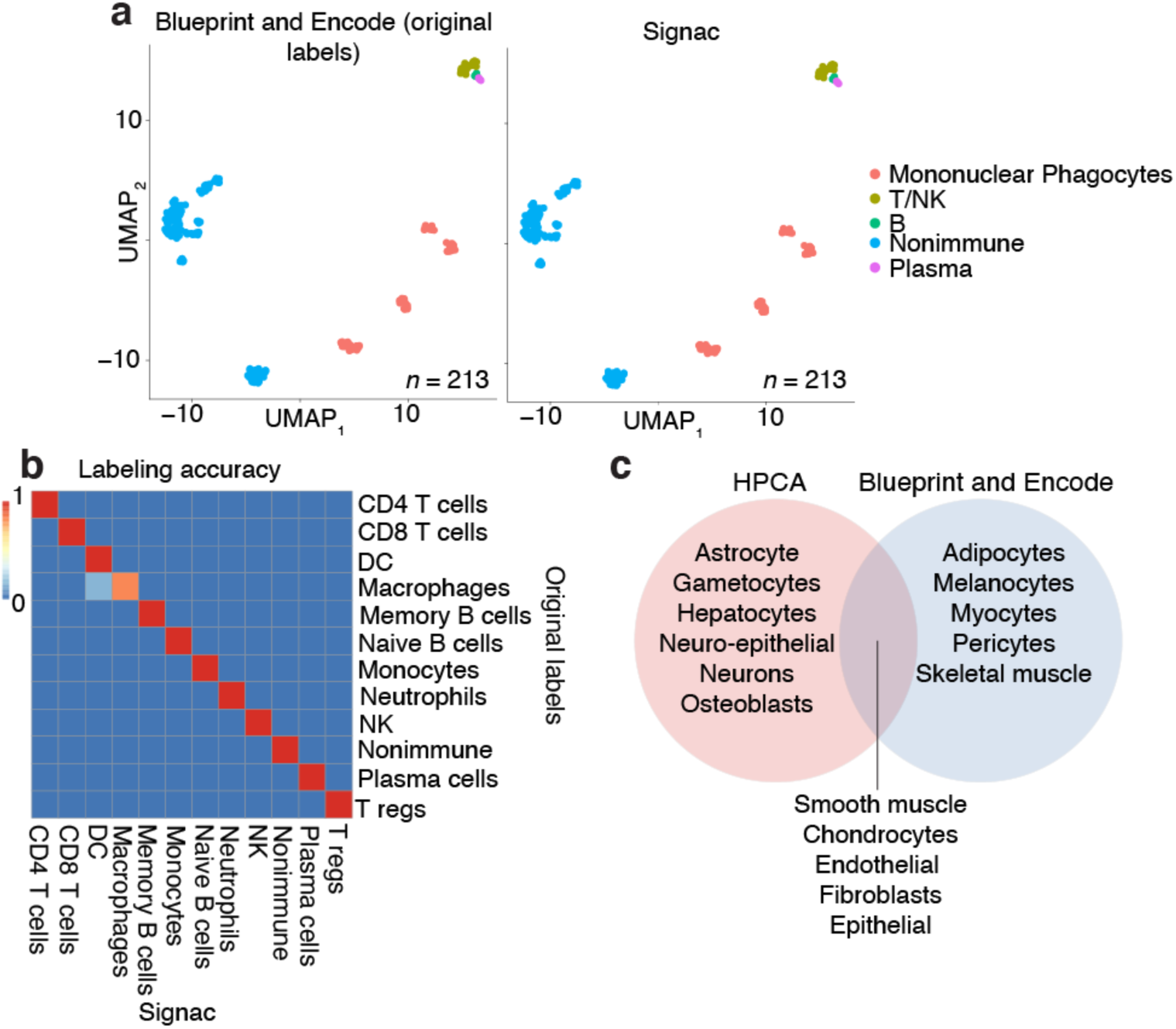
Neural networks robustly classified validation data from the Blueprint and Encode consortia. **A, UMAP plots show that cellular phenotypes were accurately classified by Signac**. In the scatter plot (left), each dot is a sample of pure cell type gene expression data that was amended labels for cellular phenotypes (colors; see legend). The data represented here are in a two-dimensional embedding (axes), in which distances correspond to transcriptional similarities between samples (closer samples are more similar); we determined this embedding with UMAP. Cellular phenotype labels (colors) were established either by the Blueprint and Encode consortium (left) with empirical measurements or by our computational approach (right). **B, Heatmap shows that distinct immune cell phenotypes were accurately classified by Signac**. Heatmap displays the fraction of the samples within each cellular phenotype category (axes) that were accurately classified by our approach (scale bar; red is more accurate; blue is less accurate). **C, Venn diagram shows that Signac accurately identified the transcriptomes of nonimmune cellular phenotypes despite never being trained to recognize them**. Venn diagram depicts the nonimmune cellular phenotypes that were specific to the HPCA reference data (left; red), that were shared between the HPCA reference and the Blueprint and Encode data (middle; purple), and that were distinct to the Blueprint and Encode data (right; blue).

**Supplemental Figure 3:**
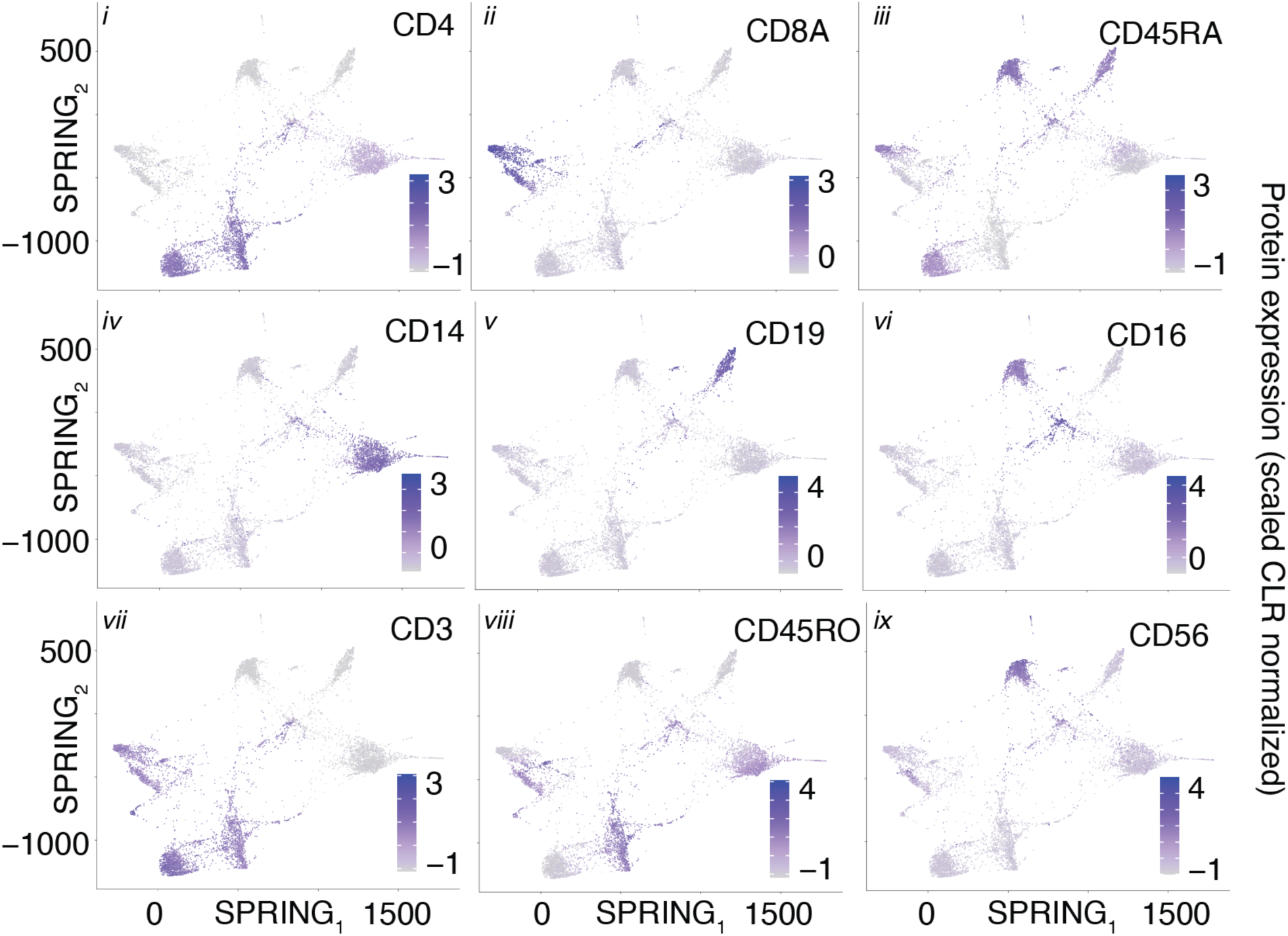
Protein expression SPRING plots validated the cellular phenotypes classified by Signac for the CITE-seq PBMCs data. Each SPRING plot (*i*-*ix*) displays the z-score transformed CLR normalized protein expression (colors) generated for each individual cell.

**Supplemental Figure 4:**
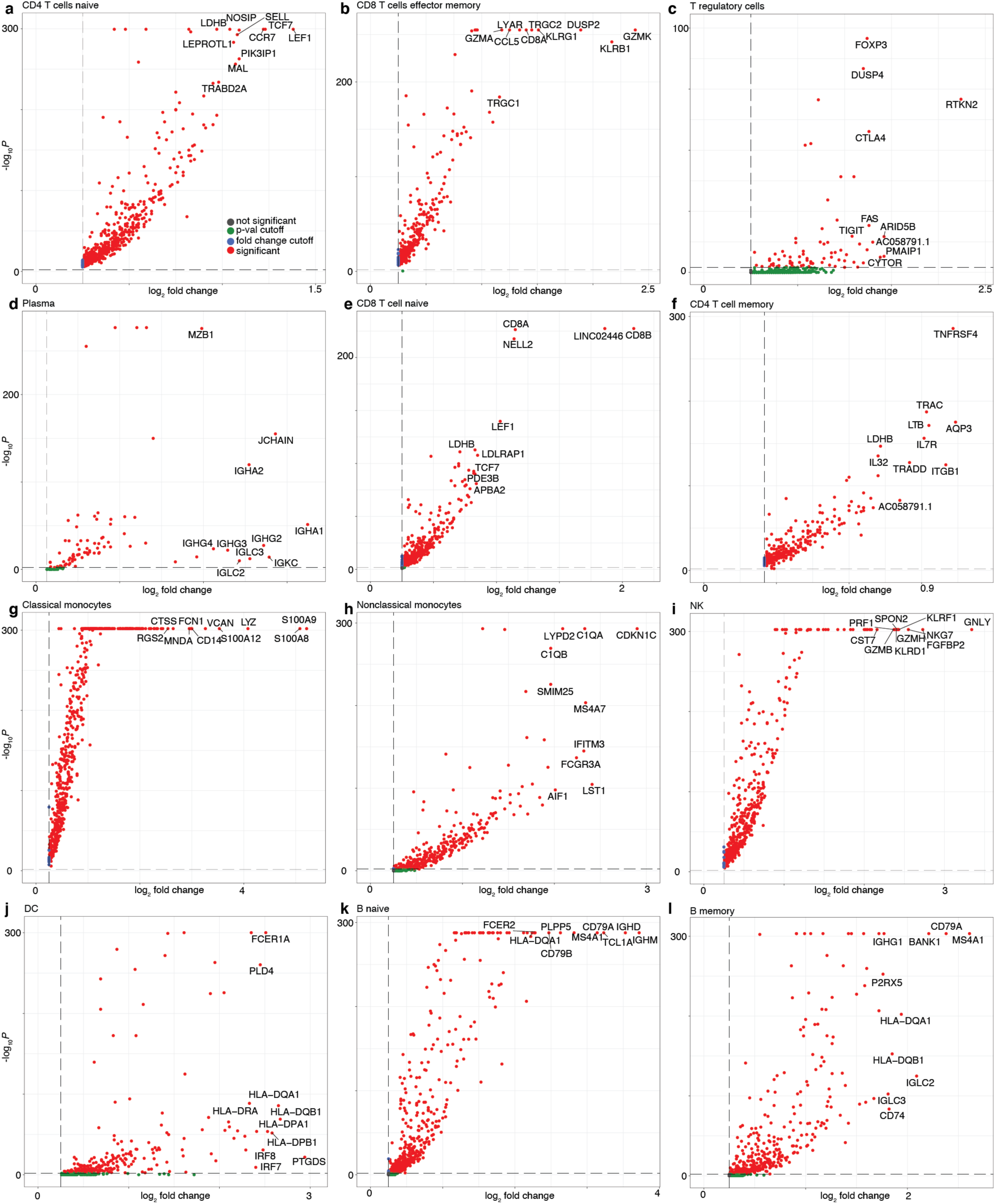
IMAGES identified from immune phenotypes in CITE-seq PBMCs single cell transcriptomes. **A-I, Volcano plots demonstrate the IMAGES identified in each cell population**. Each scatter plot depicts the statistical association (y-axis) and the average fold-change of IMAGES (each dot is a unique gene) for immune cell phenotypes. Colors (red) indicate IMAGES that passed the thresholds applied to the fold-change and adjusted p-values.

**Supplemental Figure 5:**
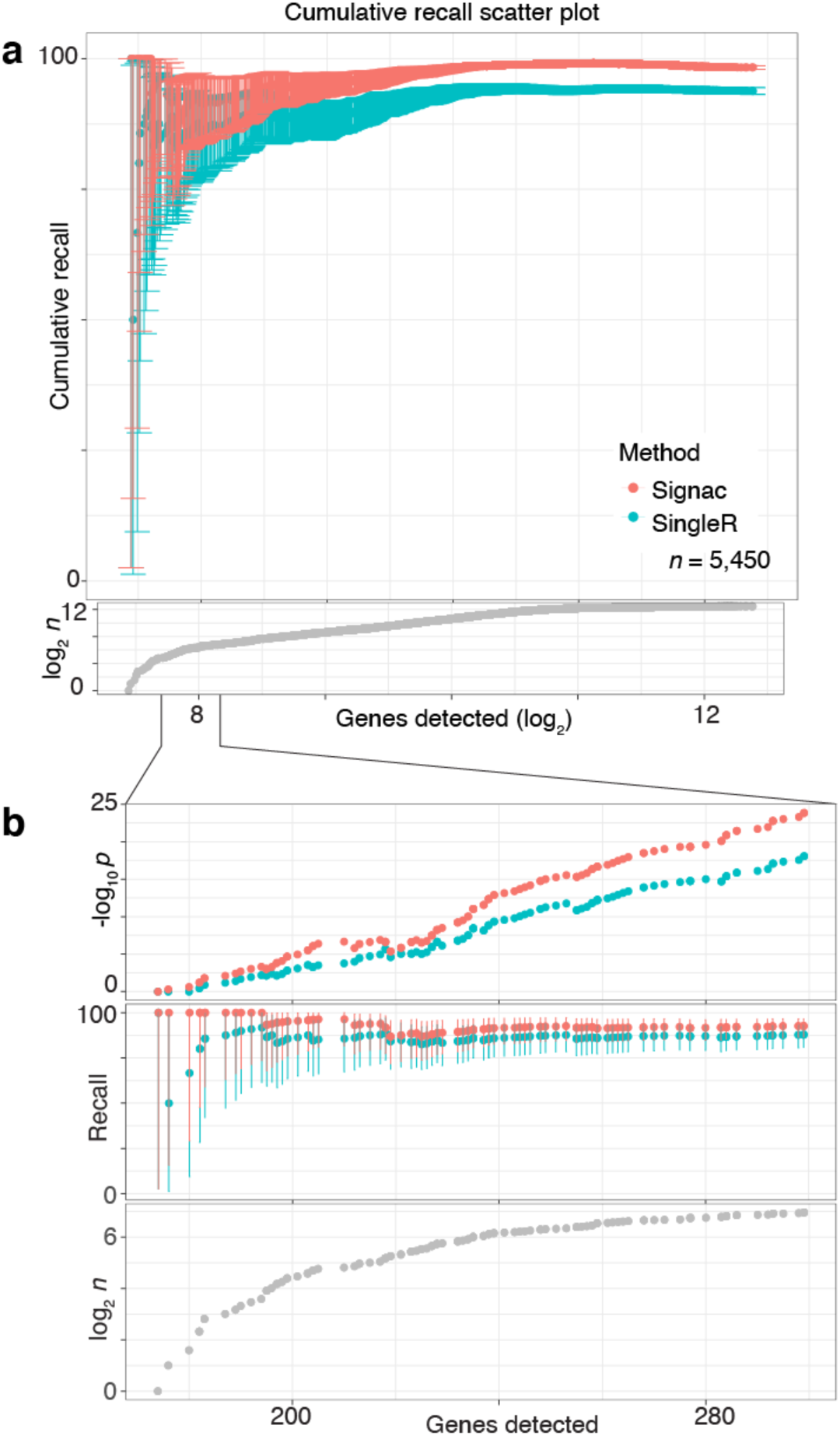
Signac accurately recalled flow cytometry labels with few genes detected. **A, Cumulative recall scatter plot showed that Signac outperformed SingleR as a function of genes detected in immune cell transcriptomes**. Plot depicts the cumulative recall (y-axis) of immune cell type labels. The labels were originally determined by flow cytometry (T cell, B cell or monocyte). Recall of these labels was calculated cumulatively as a function of the number of genes detected (x-axis) by either Signac (red) or SingleR (teal). Fibroblasts were omitted from this analysis due to broad misclassification by SingleR. Scatter plot, bottom depicts the number of single cell transcriptomes (*n*) as a function of genes detected. Error bars (top) are 95% C.I.s determined by two-sided binomial testing. **B, Inset shows stronger Signac performance at low genes detected**. Scatter plot (top) depicts the *p*-value for the two-sided binomial test and showed that Signac (red) outperformed SingleR (teal) at low sequencing depths. Scatter plots (middle; bottom) are close-ups of **Supplemental Figure 5A** for data with less than 300 genes detected.

**Supplemental Figure 6:**
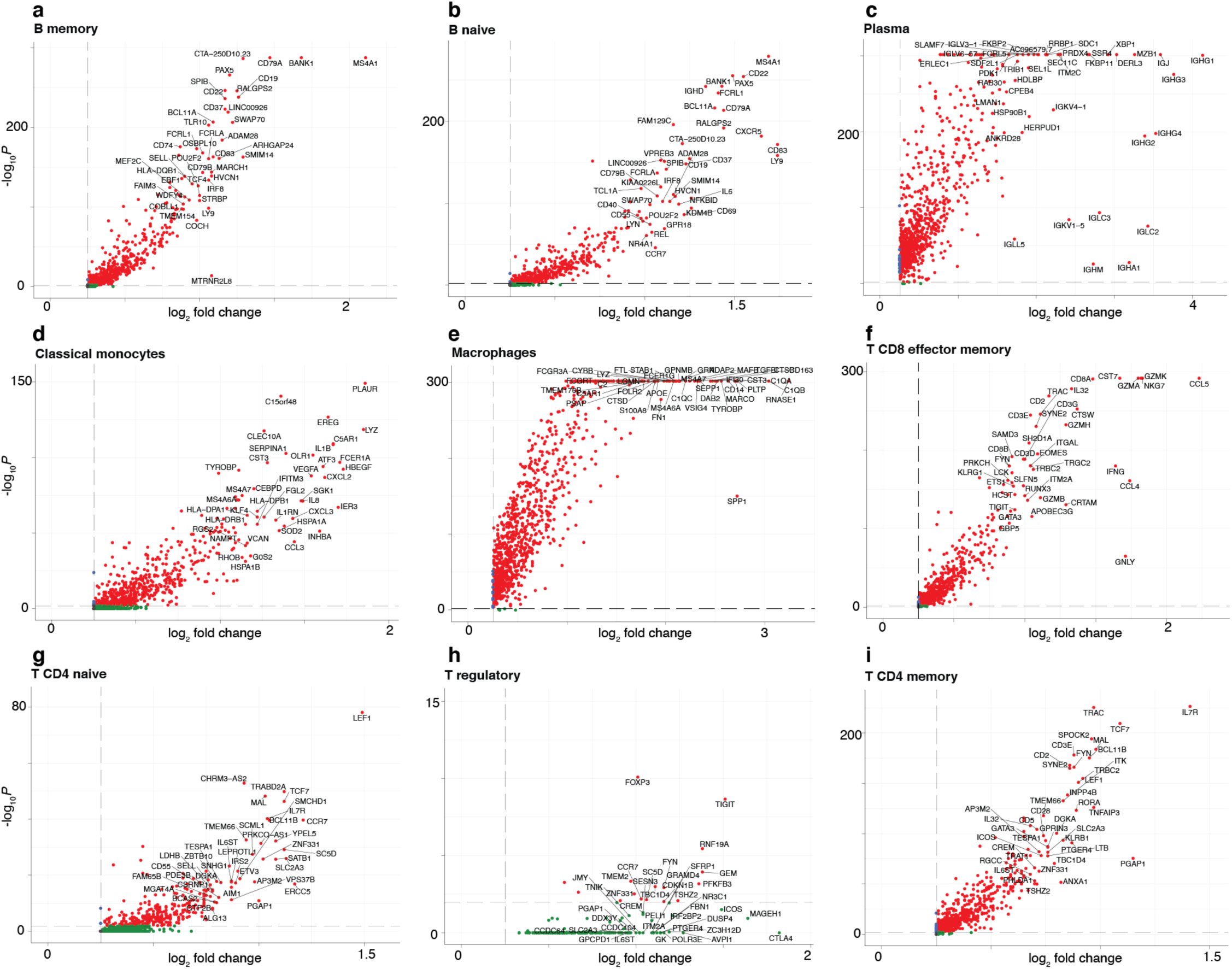
IMAGES identified from immune phenotypes in synovium single cell transcriptomes. **A-I, Volcano plots demonstrate the IMAGES identified in each cell population**. Each scatter plot depicts the statistical association (y-axis) and the average fold-change of IMAGES (each dot is a unique gene) for immune cell phenotypes. Colors (red) indicate IMAGES that passed the thresholds applied to the fold-change and adjusted p-values.

**Supplemental Figure 7:**
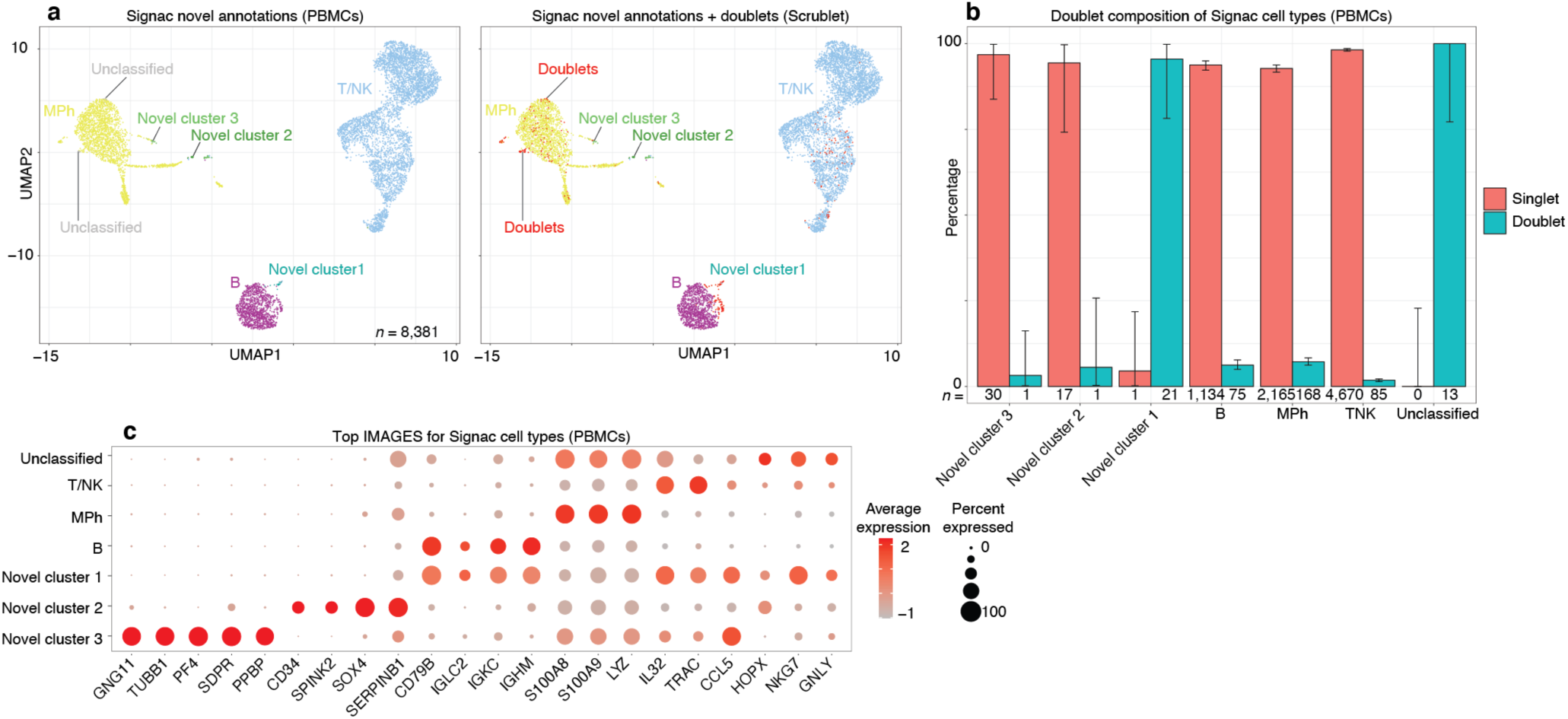
Signac-generated “unclassified” PBMCs were all identified as doublets (Scrublet), novel and classified cell type populations were mostly singlets. **A-I, Volcano plots demonstrate the IMAGES identified in each cell population. A, UMAP plot displays the cell type annotations of Signac for PBMCs (left), and the same data but with doublets classified by Scrublet (right)**. Each cell barcode (*n* = 8,381 from one human donor) was classified by Signac (left), and then doublet labels were amended to each cell with Scrublet (right; red); see <u>Methods: Signac classification</u>. **B, Bar plot reveals that unclassified cells were entirely composed of doublets, whereas novel cell populations and classified cells were mostly composed of singlets**. Each bar shows the percentage (y-axis) of each cell type (x-axis) that is a doublet (teal) or a singlet (red). Error bars correspond to 95% confidence intervals, two-sided binomial test. **C, IMAGE expression dot plot shows that unclassified cells and novel cluster 2 were doublet-like, whereas novel cluster 2 and 3 were singlet-like and enriched for known platelet and hematopoietic stem-cell gene markers**. Dot plot shows the percentage (size) of single-cell transcriptomes within a cell type (y-axis) for which non-zero expression of marker genes was observed (x-axis). Color displays the average gene expression (red indicates more expression) in each cell type category. Novel cluster 1 was enriched for IMAGES that were typically enriched in either B cells or T cells, but not both (i.e., these cells expressed both *CD79B* and *TRAC*), consistent with the view that these cells were doublets. Novel cluster 3 images (GNG11^+^ TUBB1^+^) suggested platelet-like cells, and novel cluster 2 images (CD34^+^ SPINK2^+^) suggested hematopoietic stem cell-like cells.

**Supplemental Figure 8:**
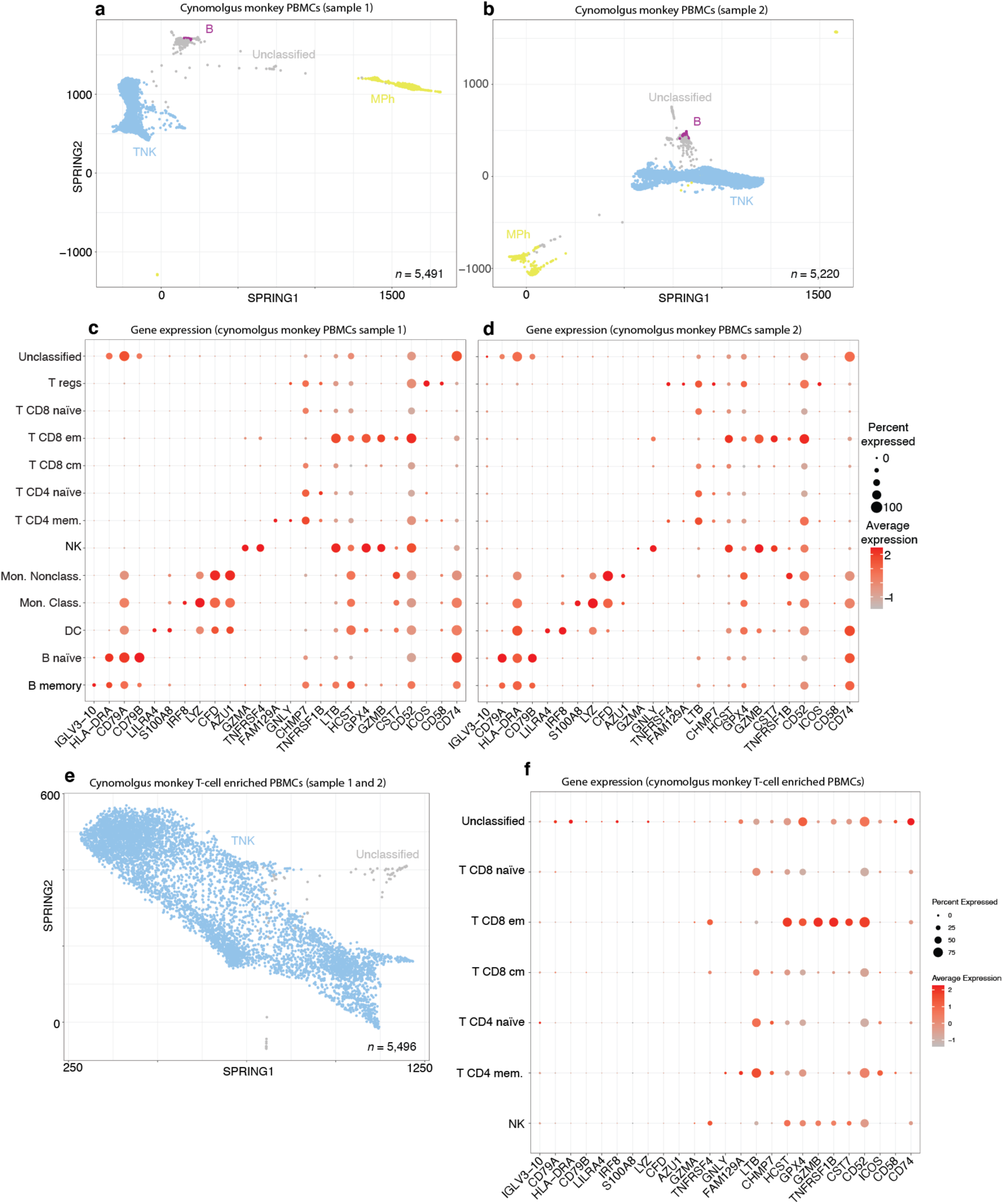
Signac accurately classified primate PBMCs without any species-specific training. **A, SPRING plot for PBMCs from cynomolgus monkey donor 3003. B, SPRING plot for PBMCs from cynomolgus monkey donor 3004. C-D, IMAGE expression dot plot shows that Signac classifications for immune phenotypes were consistent with known gene markers. E, SPRING plot for PBMCs from cynomolgus monkey samples that were enriched for T cells during sequencing.** See <u>Methods: Cross-species classification of single cell data from cynomolgus monkey PBMCs with human reference data</u>. **F, IMAGE expression dot plot shows that Signac classifications for immune phenotypes were consistent with T cell type enrichment assay**.

